# Nicotinic acid mononucleotide is an allosteric SARM1 inhibitor promoting axonal protection

**DOI:** 10.1101/2021.07.15.452560

**Authors:** Yo Sasaki, Jian Zhu, Yun Shi, Weixi Gu, Bostjan Kobe, Thomas Ve, Aaron DiAntonio, Jeffrey Milbrandt

## Abstract

SARM1 is an inducible NAD^+^ hydrolase that is the central executioner of pathological axon loss. Recently, we elucidated the molecular mechanism of SARM1 activation, demonstrating that SARM1 is a metabolic sensor regulated by the levels of NAD^+^ and its precursor, nicotinamide mononucleotide (NMN), via their competitive binding to an allosteric site within the SARM1 N-terminal ARM domain. In healthy neurons with abundant NAD^+^, binding of NAD^+^ blocks access of NMN to this allosteric site. However, with injury or disease the levels of the NAD^+^ biosynthetic enzyme NMNAT2 drop, increasing the NMN/ NAD^+^ ratio and thereby promoting NMN binding to the SARM1 allosteric site, which in turn induces a conformational change activating the SARM1 NAD^+^ hydrolase. Hence, NAD^+^ metabolites both regulate the activation of SARM1 and, in turn, are regulated by the SARM1 NAD^+^ hydrolase. This dual upstream and downstream role for NAD^+^ metabolites in SARM1 function has hindered mechanistic understanding of axoprotective mechanisms that manipulate the NAD^+^ metabolome. Here we reevaluate two methods that potently block axon degeneration via modulation of NAD^+^ related metabolites, 1) the administration of the NMN biosynthesis inhibitor FK866 in conjunction with the NAD^+^ precursor nicotinic acid riboside (NaR) and 2) the neuronal expression of the bacterial enzyme NMN deamidase. We find that these approaches not only lead to a decrease in the levels of the SARM1 activator NMN, but also an increase in the levels of the NAD^+^ precursor nicotinic acid mononucleotide (NaMN). We show that NaMN inhibits SARM1 activation, and demonstrate that this NaMN-mediated inhibition is important for the long-term axon protection induced by these treatments. Analysis of the NaMN-ARM domain co-crystal structure shows that NaMN competes with NMN for binding to the SARM1 allosteric site and promotes the open, autoinhibited configuration of SARM1 ARM domain. Together, these results demonstrate that the SARM1 allosteric pocket can bind a diverse set of metabolites including NMN, NAD^+^, and NaMN to monitor cellular NAD^+^ homeostasis and regulate SARM1 NAD^+^ hydrolase activity. The relative promiscuity of the allosteric site may enable the development of potent pharmacological inhibitors of SARM1 activation for the treatment of neurodegenerative disorders.

**Highlights:**

- NaMN binds to SARM1 N-terminal allosteric site and inhibits SARM1 NAD+ hydrolase
- NaMN inhibits SARM1 activation by stabilizing its open, inactive structure
- NMN deamidase promotes strong axonal protection by reducing NMN and increasing NaMN

## Introduction

Axon degeneration is a neuropathological consequence of both acute neuronal injury and neurodegenerative disease. SARM1 is the central executioner of axon degeneration, and a key driver of pathology in models of traumatic nerve injury (Gerdts et al., 2013; Osterloh et al., 2012), chemotherapy induced peripheral neuropathy (Cetinkaya-Fisgin et al., 2020; Geisler et al., 2019; 2016; Turkiew et al., 2017), traumatic brain injury (Henninger et al., 2016; Marion et al., 2019) and glaucoma (Ko et al., 2020). SARM1 can also induce neuronal cell death (Summers et al., 2014), and promotes photoreceptor loss in models of retinitis pigmentosa (Ozaki et al., 2020) and Leber congenital amaurosis (Sasaki et al., 2020). In addition, we and others identified human genetic variants in patients with amyotrophic lateral sclerosis that render SARM1 constitutively active, suggesting that SARM1 variants may be a risk factor for this devastating neurodegenerative disease (Bloom et al., 2021; Gilley et al., 2021). The identification of SARM1 as a therapeutic target for this wide range of neurodegenerative disorders has motivated detailed studies of the SARM1 mechanism of action in hopes of identifying novel therapeutics (Figley and DiAntonio, 2020; Krauss et al., 2020).

SARM1 is the founding member of the TIR-domain family of NAD^+^ hydrolases (Essuman et al., 2017; 2018; Horsefield et al., 2019; Wan et al., 2019). The SARM1 protein comprises an N-terminal Armadillo repeat motif (ARM) autoinhibitory domain, SAM domains required for oligomerization, and a C-terminal TIR domain that catalyzes NAD^+^ hydrolysis (Essuman et al., 2017; Sporny et al., 2020). SARM1 forms an octamer (Horsefield et al., 2019; Sporny et al., 2019), and interactions across multiple domain interfaces both within and across subunits are required to keep the SARM1 octamer inactive (Shen et al., 2021). Upon injury, the autoinhibitory N-terminus releases the TIR domain to activate the NAD^+^ hydrolase. Interestingly, SARM1 not only acts upon NAD^+^, but its activation state is also regulated by NAD^+^ and the NAD^+^ precursor nicotinamide mononucleotide (NMN) (Figley et al., 2021; Jiang et al., 2020; Sporny et al., 2020; Zhao et al., 2019). In the off-state, NAD^+^ binds to an allosteric site in the autoinhibitory ARM domain (Jiang et al., 2020; Sporny et al., 2020). Upon axon injury, levels of the NAD^+^ biosynthetic enzyme NMNAT2 fall, thereby leading to a drop in the levels of NAD^+^ and an increase in the levels of NMN (Di Stefano et al., 2015; Gilley and Coleman, 2010). NMN binds the allosteric site much more tightly than does NAD^+^, and so when the NMN/ NAD^+^ ratio rises, NMN displaces NAD^+^ from the allosteric site, initiating a conformational change that activates the NAD^+^ hydrolase (Figley et al., 2021). This understanding of the SARM1 activation mechanism explains prior findings such as 1) overexpression of the NAD^+^ synthesis enzyme NMNAT1 inhibits SARM1 activation (Sasaki et al., 2016) and 2) inhibition of NAMPT, the NMN synthesizing enzyme, reduces cellular NMN levels and protects axons from degeneration (Di Stefano et al., 2015; Sasaki et al., 2009). However, there are some prior findings that are harder to understand. Why, for example, does reduction of NMN due to treatment with the NAMPT inhibitor FK866 only lead to short-term axonal protection (Di Stefano et al., 2015; Sasaki et al., 2009), while expression of *E. coli* NMN deamidase, which also reduces NMN levels, provides dramatically longer-term axonal protection (Di Stefano et al., 2015; 2017)? Moreover, how is the short-term protection afforded by FK866 transformed into long-lasting axon preservation by the addition of NaR, a NAMPT-independent NAD^+^ precursor that bypasses NMN production (Liu et al., 2018)? Interestingly, both scenarios with profound axon protection boost the levels of deamidated NAD^+^ precursors.

NAD^+^ biosynthesis can occur along two parallel tracks. The salvage pathway works on amidated molecules, with NAMPT converting nicotinamide to NMN and NMNAT enzymes then converting NMN to NAD^+^. The Preiss Handler pathway works on deamidated molecules, with nicotinic acid phosphoribosyl transferase (NaPRT) converting nicotinic acid to nicotinic acid mononucleotide (NaMN) and NMNAT enzymes then converting NaMN to nicotinic acid adenine dinucleotide (NaAD), with NAD^+^ synthetase finally generating NAD^+^ from NaAD. The two axoprotective treatments described above both lead to the production of NaMN. The enzyme NMN deamidase directly converts NMN to NaMN, while the precursor NaR is converted to NaMN by the cellular enzymes NRK1 and NRK2 (nicotinamide riboside kinases 1&2) (Belenky et al., 2009; Gong et al., 2013). This observation led us to test the hypothesis that NaMN, or another component of the deamidated pathway, may inhibit SARM1 and thereby contribute to the strong axon protection induced by these treatments. We demonstrate that the addition of NaR to FK866 treated axons post-axotomy promotes strong axonal protection without boosting NAD^+^ biosynthesis. Instead, NaMN derived from NaR binds the SARM1 ARM domain and inhibits the activation of the SARM1 NAD^+^ hydrolase. Likewise, we show that the strong axonal protection afforded by neuronal expression of *E. coli* NMN deamidase is mediated not only by an NMN decrease, but also by an NaMN increase. These results demonstrate that the SARM1 allosteric pocket binds diverse components of the NAD^+^ metabolome to monitor cellular NAD^+^ homeostasis and regulate SARM1 NAD^+^ hydrolase activity. The identification of NaMN as a SARM1 inhibitor may facilitate the development of novel approaches for blocking SARM1 activation as treatments for neurodegenerative disorders.

## Results

The application of nicotinic acid riboside (NaR) and FK866, an inhibitor of NAMPT that synthesizes NMN, strongly delays vincristine-induced axon degeneration (Liu et al., 2018). To test the effect of the co-administration of NaR and FK866 on axon degeneration after axotomy, dorsal root ganglion (DRG) neurons were treated with various concentrations of FK866 and NaR pre- and post-axotomy. As previously reported (Di Stefano et al., 2015; Sasaki et al., 2016; 2009), the pretreatment of FK866 (100 nM) for 24 hours before axotomy provided short-term (6 hours) axon protection (Figure 1A, B). In contrast, in FK866 pretreated axons, NaR (100 μM) addition to the culture medium at 10 minutes post-axotomy conferred prolonged axonal protection (24 hours) (Figure 1A, B). This effect is dose dependent, with an EC_50_ of ∼18 μM NaR in the culture media (Figure 1E). NaR alone provides no axonal protection, demonstrating that combination treatment is required for the long-term axon preservation. Moreover, this effect is selective for NaR, as FK866 preincubation with nicotinic acid, another deamidated NAD^+^ precursor, provided no additional protection to FK866 preincubation. We have previously demonstrated that inhibiting SARM1 function as much as three hours after injury is sufficient to block axon degeneration (Gerdts et al., 2015; Sasaki and Milbrandt, 2010). We varied the timing of NaR application in conjunction with FK866 pretreatment and found that NaR addition at 10 minutes, two, or three hours post-axotomy provided indistinguishable and potent axonal protection lasting at least 72 hours (Figure 1C, D, F). It is notable that NaR potently protects axons when added three hours post-axotomy, as by this time NMNAT2, the major NMNAT enzyme in the axon, is largely degraded (Gilley and Coleman, 2010; Summers et al., 2019). A prior study demonstrating that a combination treatment with FK866 and NaR could protect axons from vincristine treatment, suggested that the role of NaR was to boost NAD^+^ levels (Liu et al., 2018); however, our finding that NaR can be added after the NAD^+^ biosynthetic pathway is impaired calls this NAD^+^ synthesis hypothesis into questions.

**Figure 1.**
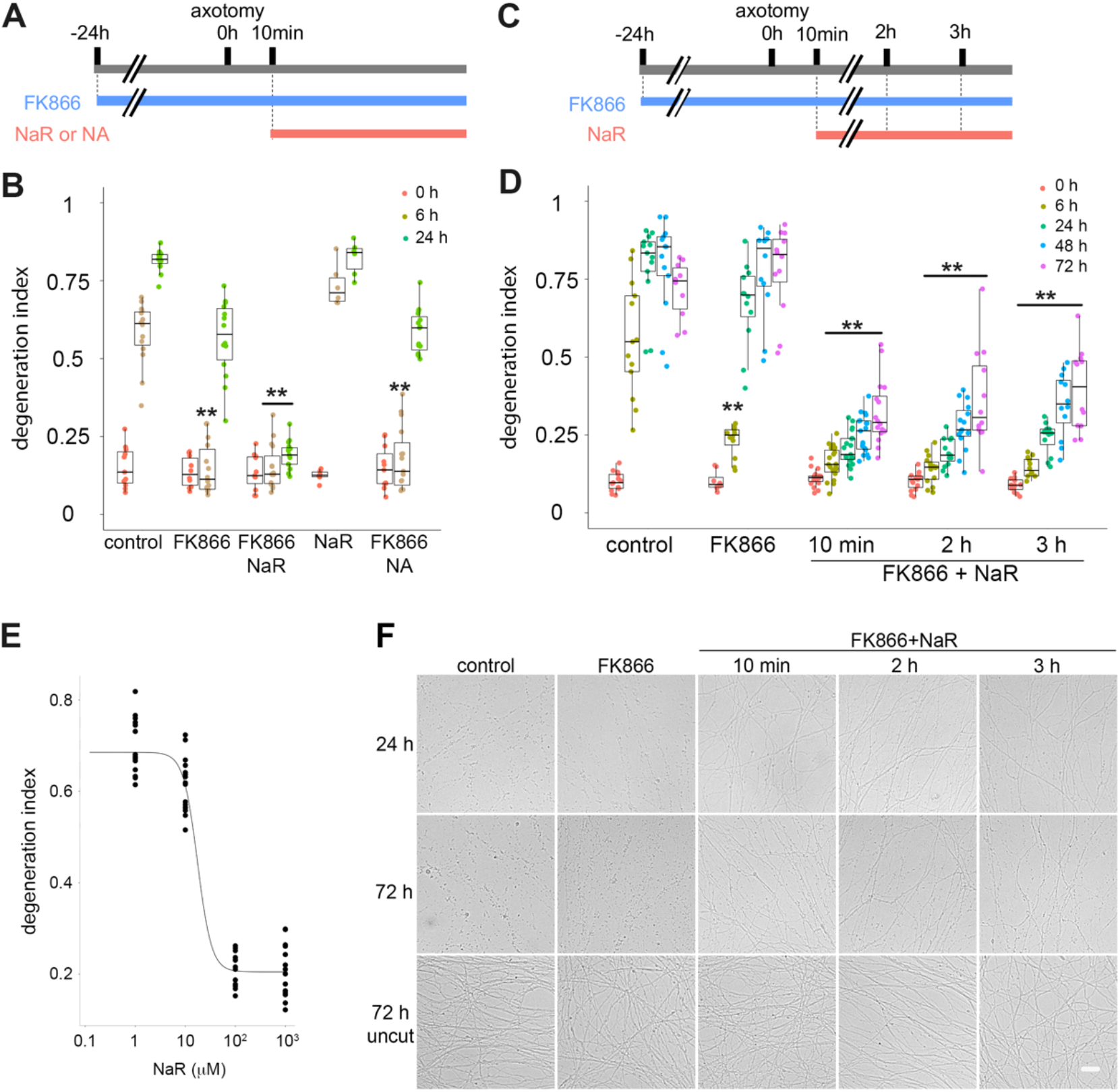
NaR provides prolonged axonal protection after axotomy in the presence of FK866. A) Time scheme showing the timing of axotomy, chemical addition (FK866 100nM, NaR or NA 100 μM), and axon degeneration assay used in (B). B) Axonal degeneration 0 to 24 h post-axotomy with treatments described in (A) was quantified. Statistical analysis was performed by two-way ANOVA with Tukey post-hoc test. F(4, 164)=178.4, p<1.0×10^−16^ among groups including control, FK866 (100 nM), FK866 (100 nM)+NaR (100 μM), NaR (100 μM), and FK866+nicotinic acid (NA). dex of control at the same time post-axotomy. C) Time scheme showing the timing of axotomy, chemical addition (FK866 100nM, NaR 100 μM), and axon degeneration assay used in (D). D) Axonal degeneration 0 to 72 h post-axotomy with treatments described in (C) was quantified. While control or NaR-treated axons degenerate within 6 h after axotomy, FK866 showed axonal protection at 6 h, but not 24 h. NaR addition 10 min to 3 h post axotomy provided strong axonal protection up to 72 h. Statistical analysis was performed by two-way ANOVA with a Tukey post-hoc test. F(4, 317)=269.1, p<1.0×10^−16^ among groups including control, FK866, FK866+NaR (10 min, 2h, 3h). **p<1 × 10^−5^ denotes the significant difference compared with control 0 h post-axotomy. E) Dose response curve showing NaR-mediated axonal protection in the presence of 100 nM FK866. EC_50_ = 18 μM. F) Representative images of DRG axons at 24 h or 72 h after axotomy or 72 h without injury (uncut) with the indicated treatments. Scale bar, 20 μm.

To investigate the mechanism of NaR mediated axonal protection, we first explored the hypothesis that NaR functions through enhanced NAD^+^ synthesis (Liu et al., 2018). For NaR to promote axonal NAD^+^ synthesis, it must be converted to NaMN by either NRK1 or NRK2, then to NaAD by NMNAT2, and finally to NAD^+^ by NAD^+^ synthetase (NADSYN) (Liu et al., 2018) (Figure 2A). However, the short-lived NMNAT2 enzyme is quickly lost after either axotomy or vincristine treatment (Geisler et al., 2019; Gilley and Coleman, 2010; Summers et al., 2019). We first asked whether NaR or FK866 influenced the level of axonal NMNAT2 in injured axons. Neurons were incubated with FK866 for 24 hours before axotomy, then NaR was added 10 minutes after injury (Figure 2B). We used western blotting to monitor NMNAT2 levels and found that NMNAT2 was reduced more than four-fold within three hours under all conditions tested (Figure 2C), indicating that NaR and FK866 inhibition of axon degeneration is not mediated by increased NMNAT2 stability. We next analyzed the effects of NaR and FK866 on axonal NAD^+^ levels before and after axotomy. Neurons were incubated with FK866 for 24 hours before axotomy, then NaR was added three hours after injury, a time at which it is potently protective (Figure 2D). Axonal metabolites were collected at 0 or 27 hours after axotomy and measured using LC-MSMS (Figure 2E). Consistent with previous reports (Sasaki et al., 2016; 2009), basal axonal NAD^+^ levels were dramatically reduced by FK866 treatment (Figure 2E, panel NAD^+^, 0h control vs 0h FK866) and axotomy (Figure 2E, panel NAD^+^, 0 h control vs 27h control). While axonal NaMN was not increased in neurons treated by NaR alone likely due to their degenerating condition, the addition of NaR together with FK866 significantly increased NaMN and NaAD in transected axons (Figure 2E) and prevented their degeneration (Figure 1D); however, it did not rescue axotomy-induced or FK866-mediated NAD^+^ depletion. Hence, NaR is not conferring protection by increasing NAD^+^ levels, and so must work through a different mechanism.

**Figure 2.**
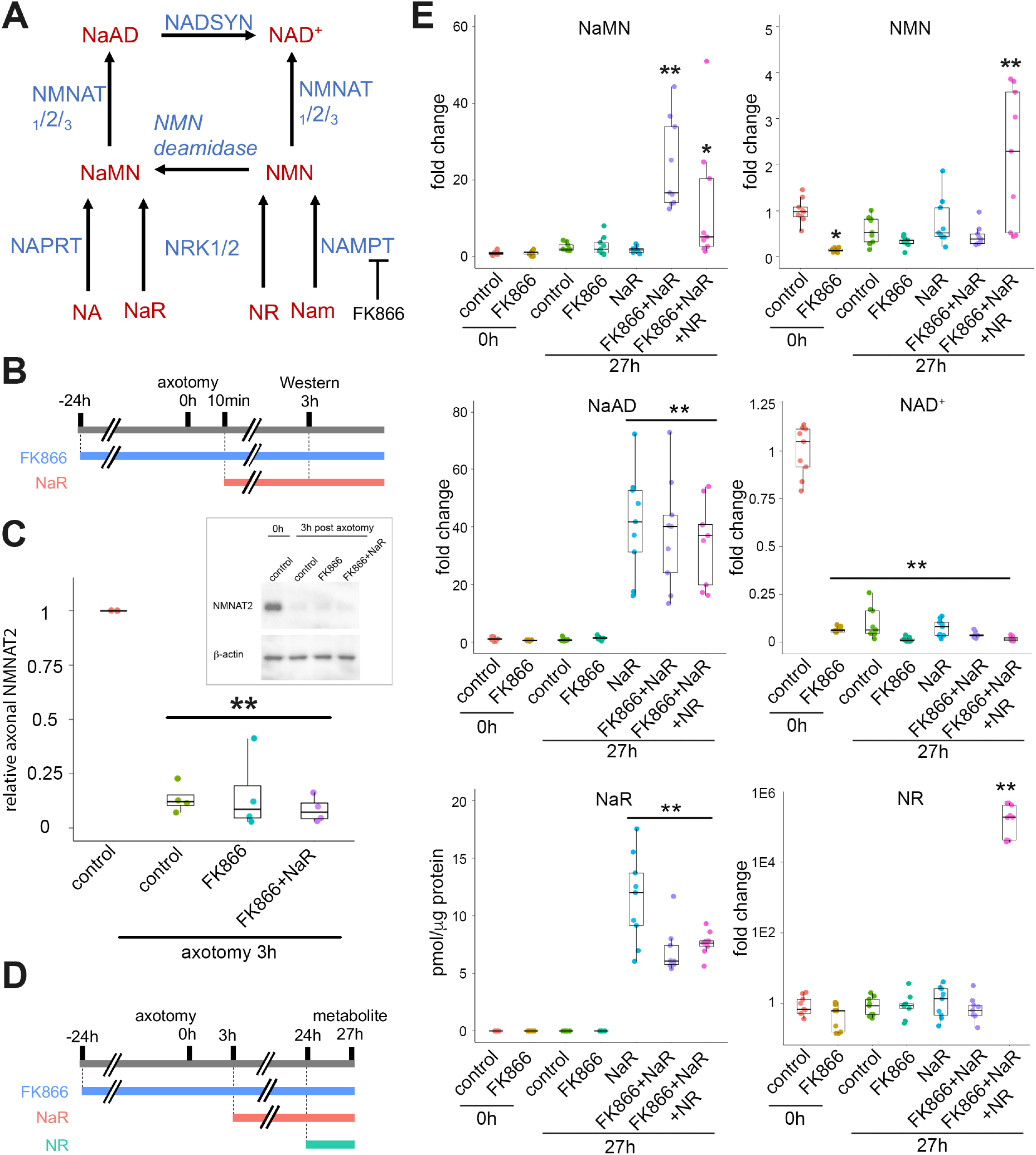
NaR-mediated NaMN and NaAD increases do not support NAD^+^ synthesis in transected axons. A) Diagram of mammalian nicotinamide adenine dinucleotide (NAD^+^) biosynthesis pathways. Nam; nicotinamide, NA; nicotinic acid, NMN; nicotinamide mononucleotide, NR; nicotinamide riboside, NaR; nicotinic acid riboside, NaMN; nicotinic acid mononucleotide, NAD^+^; nicotinamide adenine dinucleotide, NaAD; nicotinic acid adenine dinucleotide. FK866 is an inhibitor of NAMPT. B) Time scheme showing the timing of axotomy, chemical addition (FK866 100nM, NaR100 μM), and western blotting used in (C). C) Axonal NMNAT2 normalized by actin at 0 or 3 h post-axotomy are shown. Data were normalized to the control axons 0 h post-axotomy. Statistical analysis was performed by one-way ANOVA with a Tukey post-hoc test. F(3, 11)=57.6, p=1.5×10^−6^ among groups. **p<5.1×10^−7^ denotes the significant difference compared with the amount of NMNAT2 in control axons at 0 h post-axotomy. Inset shows a representative western blot, showing axonal NMNAT2 0 or 3 h post-axotomy with indicated conditions. D) Time scheme showing the timing of axotomy, chemical addition (FK866 100nM, NaR 100 μM, NR 100 μM), and metabolite measurement used in (E). E) Axonal metabolite analyses 0 and 27 h post-axotomy with treatments described in (D) are shown. NaR in control axons was below the lower limit of detection. Note that metabolites post 27 hours axotomy were measured from degenerating axons in control, FK866, NaR, or FK866+NaR+NR. Statistical analysis was performed by one-way ANOVA with Benjamini-Hochberg post-hoc test. F(6, 56)=11.3, p=3×10^−8^ for NMN, F(6, 56)=10.4, p=1.1×10^−7^ for NaMN, F(6, 56)=27.7, p=4.4×10^−15^ for NaAD, F(6, 56)=311.7, p<2×10^−16^ for NAD^+^, F(6, 56)=74.1, p<2×10^−16^ for NaR, F(6, 56)=14.4, p=7.8×10^−10^ for NR. **p<1×10^−5^ and *p<0.05 denote the significant difference compared with the amounts of metabolites in control axons at 0 h post-axotomy.

Nicotinamide riboside kinases (NRK1 or 2) metabolize NaR as well as nicotinamide riboside (NR, the amidated form of NaR) in mammalian cells (Belenky et al., 2009; Gong et al., 2013). NRK1/2 phosphorylates NaR and produces NaMN. We set to determine the role of NRK enzymes in NaR-mediated axonal protection. We generated NRK1/2 double knockout mice and found they are viable and fertile without overt behavioral phenotypes (Supplemental Figure 1A). NRK1 deficient mice did not display any overt phenotype (Ratajczak et al., 2016) and our results indicated dispensable roles of NRK1 and 2 dependent processes for mouse survival. We did not test the glucose intolerance, insulin resistance, and hepatosteatosis that were reported in NRK1 deficient mice (Sambeat et al., 2019). NRK1/2 double knockout (dKO) DRG neurons were healthy with qualitatively normal axonal growth (Supplemental Figure 1B). To assess the metabolic fate of NaR, NRK1/2 dKO neurons were treated with NaR in the presence or absence of lentiviral-mediated expression of NRK1. Unlike in NaR treated wild type neurons (Figure 2E), the synthesis of NaMN and NaAD was completely blocked in NRK1/2 dKO neurons after NaR addition (100 μM for 24 hours), confirming the depletion of NRK1/2 in these neurons. This phenotype was rescued by the exogenous expression of NRK1, which stimulated high levels of NaMN and NaAD in NaR-treated neurons (Figure 3A, B). Hence, NRK1 or NRK2 are absolutely required for the conversion of NaR and NR (Supplemental Figure1C), but are not required for cellular or animal viability. Next, we measured axonal degeneration in add-back experiments in which NRK1 was re-introduced into NRK1/2 dKO neurons using lentivirus. The addition of NaR 10 minutes post-axotomy to FK866-treated wild type neurons showed strong axonal protection. This treatment regimen was ineffective in NRK1/2 dKO axons, with robust axon degeneration in the presence of NaR and FK866 (Figure 3C, D). When NRK1 was re-expressed in these dKO neurons, NaR and FK866 joint treatment once again provided strong axonal protection. Hence, NRK-mediated NaMN synthesis from NaR is necessary for NaR to promote axon preservation.

**Figure 3.**
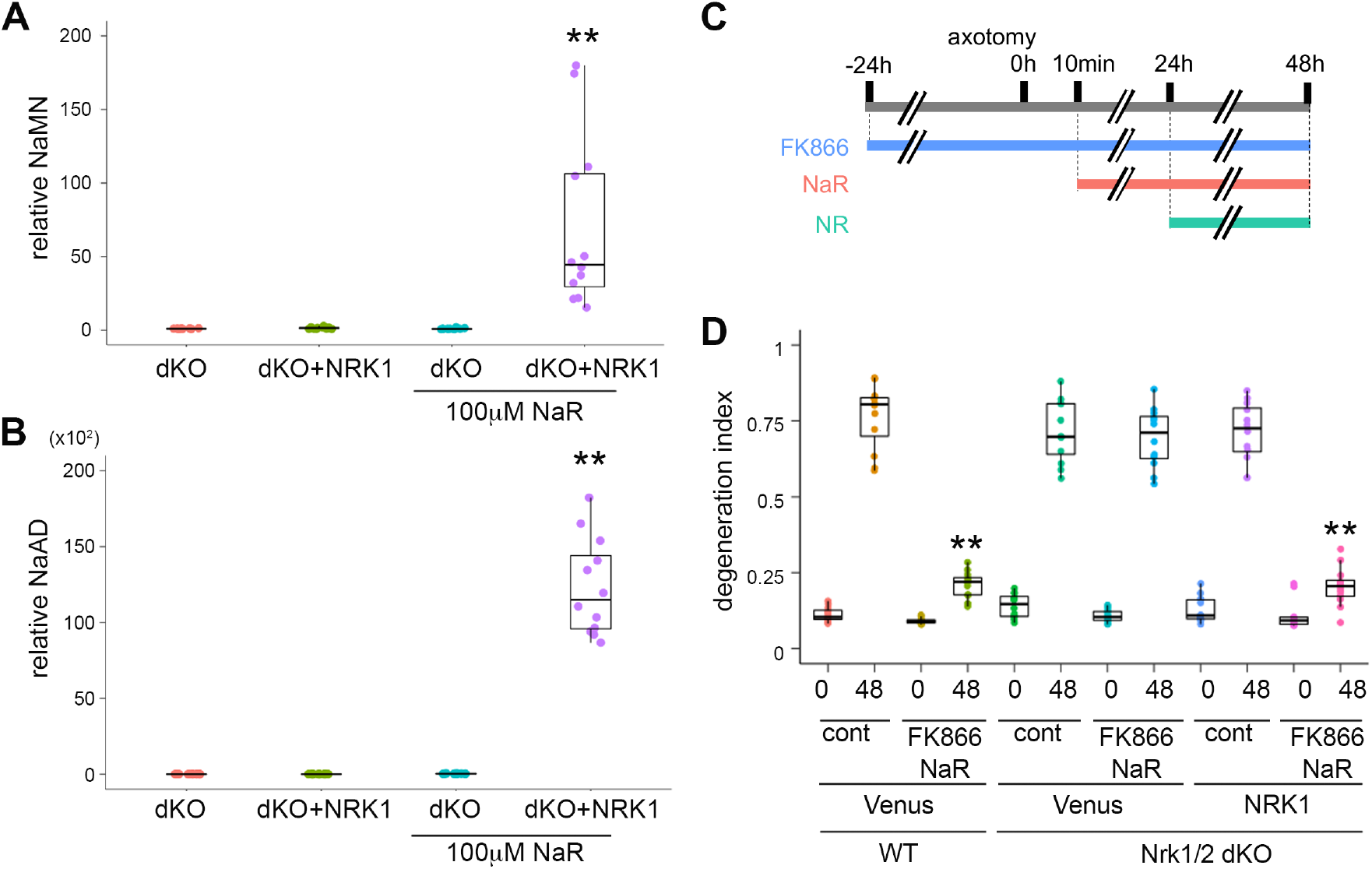
NaMN synthesis is required for axonal protection by NaR. A, B) Metabolite analysis from NRK1 and NRK2 double knockout (NRK1/2 dKO) cells expressing Venus (dKO) or mouse NRK1 (dKO+NRK1) via lentivirus. NaMN (A) or NaAD (B) were low in NRK1/2 dKO neurons with or without NaR addition (100 μM, 24 hours). Exogenously expressed NRK1 produces high NaMN and NaAD in NRK1/2 dKO neurons. Statistical analysis was performed by one-way ANOVA with a Tukey post-hoc test. F(3, 44)=16.5, p=2.5×10^−7^ for NaMN and F(3,44)=181.1 P<2 ×10^−16^ for NaAD. **p<1×10^−5^ denotes the significant difference compared with metabolites of the NRK1/2 dKO control neurons. C) Time scheme showing the timing of axotomy, chemical addition (FK866 100 nM, NaR 100 μM, NR 100 μM), and axon degeneration assay used in (D). D) Axonal degeneration 0 or 48 h post-axotomy with treatments described in (C) was quantified. Wild-type axons were protected by NaR (100 μM) with FK866 (100 nM). NRK1/2 dKO axons were not protected by NaR with FK866, however exogenous expression of NRK1 in NRK1/2 dKO neurons rescued the axonal protection by NaR with FK866. Statistical analysis was performed by two-way ANOVA with a Tukey post-hoc test. F(5, 130)=68.7, p<2.0×10^−16^ among groups. **p<1×10^−5^ denotes the significant difference compared with the degeneration index of control at the same time post-axotomy.

SARM1 activation is regulated by the balance between NMN and NAD^+^, the substrate and product of NMNAT enzymes, respectively (Figley et al., 2021). This model can explain the short-lived protection afforded by FK866 alone—at baseline, NMN is depleted, promoting axon protection, but as NMNAT2 is lost, post-axotomy, NAD^+^ levels fall, enabling even low levels of NMN to activate SARM1. However, this model cannot explain the enhanced protection provided by addition of NaR to FK866, because NaR does not rescue NAD^+^ levels (Figure 2E). Instead, NaR promotes the synthesis of NaMN (Figure 3A-D), which is structurally similar to NMN. Therefore, we hypothesize that NaMN or its derivatives compete with NMN to bind SARM1 and, hence, block NMN-mediated SARM1 activation. As a first test of this hypothesis, we increased NMN production in injured axons preserved by combining NaR and FK866 treatment. Neurons were pre-treated with FK866 for 24 hours before axotomy, and then NaR was added three hours post-axotomy. At 24 hours post-axotomy, these non-degenerating axons were treated with NR (100 μM) to increase NMN levels in a NAMPT-independent manner (Figley et al., 2021) (Figure 4A). As expected, axonal NMN was reduced by FK866 treatment (Figure 2E, control 0 h vs FK866 0 h), and these levels were similar to those in neurons treated with NaR or NaR plus FK866. In contrast, the addition of NR to neurons treated with NaR plus FK866 twenty-four hours post-axotomy resulted in large increases in NMN within three hours (Figure 2E, panel NMN). Notably, NAD^+^ was not increased in these axons likely due to the loss of NMNAT2 (Figure 2E, panel NAD^+^). We also observed a decrease of NaMN after NR treatment, potentially due to competition for the NRK enzymes that convert both NR and NaR to NMN and NaMN, respectively (Figure 2E, panel NaMN). We next assessed axonal integrity in these injured, but preserved axons, after NR addition (Figure 4A). While axons treated with combination of NaR and FK866 were intact at 48 hours post-axotomy, the addition of NR at 24 hours post-axotomy resulted in very rapid and robust axonal degeneration of these NaR and FK866 treated axons (Figure 4B, C). Hence, the high levels of NMN produced in response to NR eliminate the axon protection provided by NaR and FK866 treatment. This is consistent with the model that NaMN competes with NMN for binding to the allosteric site of SARM1.

**Figure 4.**
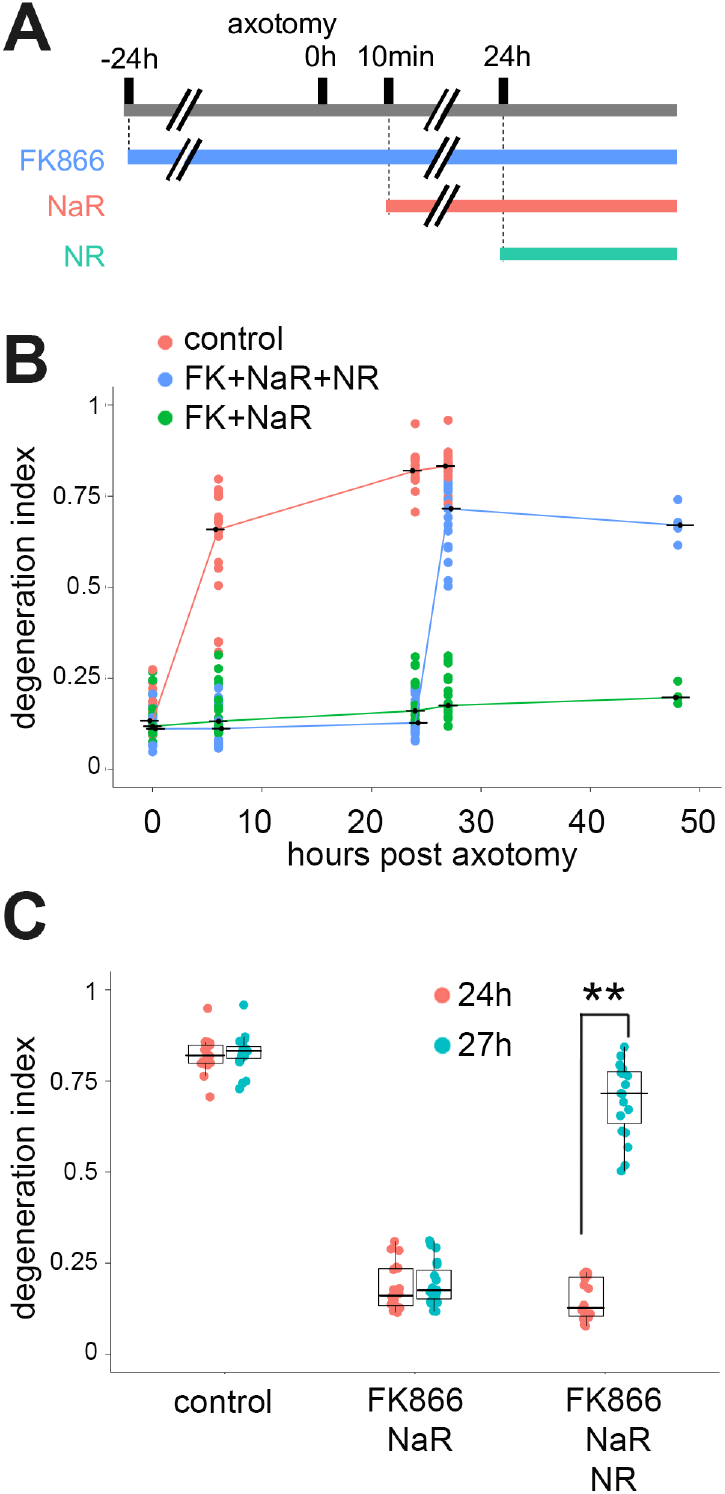
The addition of NR blocks NaR-mediated axonal protection. A) Time scheme showing the timing of axotomy, chemical addition (FK866 100 nM, NaR 100 μM, NR 100 μM), and axon degeneration assay used in (B). B) Axonal degeneration 0 to 48 h post-axotomy with treatments described in (A) was quantified. While axons treated with NaR (100 μM) and FK866 (100 nM) (FK+NaR) showed axonal protection up to 48 h, NR (100 μM) addition at 24 h post-axotomy induced axon degeneration within three hours in NaR+FK866 treated axons (FK+NaR+NR). C) Statistical analysis of axon degeneration at 24 and 27 h post axotomy in (B) was performed by two-way ANOVA with Tukey post-hoc test. F(2, 108)=924.9, p<2.0×10^−16^ among groups. **p<1×10^−5^ denotes the significant difference compared with the degeneration index of control at the same time post-axotomy.

Our metabolite and axon degeneration data support the hypothesis that NaMN or its derivatives directly inhibit SARM1 activation by competing with NMN for binding to the allosteric pocket in the autoinhibitory N-terminal ARM domain. In this model, 1) NaMN or its derivatives will directly inhibit SARM1 NAD^+^ hydrolysis activity; 2) the SARM1 N-terminal autoinhibitory domain will be required for NaMN-mediated inhibition; and 3) NMN will mitigate this inhibition. To directly test these predictions, we performed an *in-vitro* SARM1 NAD^+^ hydrolysis assay. Strep-tagged human SARM1 (hSARM1) was expressed in HEK293T cells and affinity purified and mixed with NMN, NaMN, or NaAD. The NAD^+^ hydrolase reaction was initiated by addition of NAD^+^ (10 μM) and the rate of ADPR production, monitored by HPLC, was quantified to calculate hSARM1 NAD^+^ hydrolase activity. NMN activated hSARM1 NAD^+^ hydrolase activity with a Ka of 8.7 μM (Figure 5A), which is very similar to the binding constant of NMN for the SARM1 N-terminal domain (Figley et al., 2021). We next tested the effects of NaMN and NaAD, the major metabolites derived from NaR (Figure 2E), on hSARM1 NAD^+^ hydrolase activity. We found that NaMN inhibits ADPR production in a dose-dependent manner with an IC_50_ of 93 μM in the absence of NMN, demonstrating that NaMN can directly inhibit SARM1 NAD^+^ hydrolase activity (Figure 5B). In contrast, NaAD did not inhibit hSARM1 NAD^+^ hydrolase activity (Figure 5B), suggesting that the active inhibitory component in neurons is NaMN and not NaAD. This NaMN-mediated inhibition could either be due to binding to the allosteric site at the N-terminus, or the orthosteric site in the C-terminal TIR domain NAD^+^ hydrolase. To distinguish between these two possibilities, we performed similar activity assays using a truncated version of hSARM1 lacking the N-terminal ARM domain (ΔARM-hSARM1) that houses the allosteric site. ΔARM-hSARM1 maintains enzymatic activity, but was not inhibited by NaMN (Figure 5D). Hence, NaMN does not inhibit SARM1 activity at the orthosteric site. To test whether NaMN inhibition competes with NMN-mediated activation, we assessed NaMN inhibition of full-length SARM1 in the presence of 5 μM NMN. We found that this small amount of NMN shifts the IC_50_ of NaMN by more than 50 μM to 147 μM, consistent with the much higher affinity of NMN for binding to SARM1 (Figure 5C). These results support the hypothesis that NaMN binds to the SARM1 N-terminal allosteric NMN/ NAD^+^ binding site and competes with NMN to inhibit SARM1 NAD^+^ hydrolase activity.

**Figure 5.**
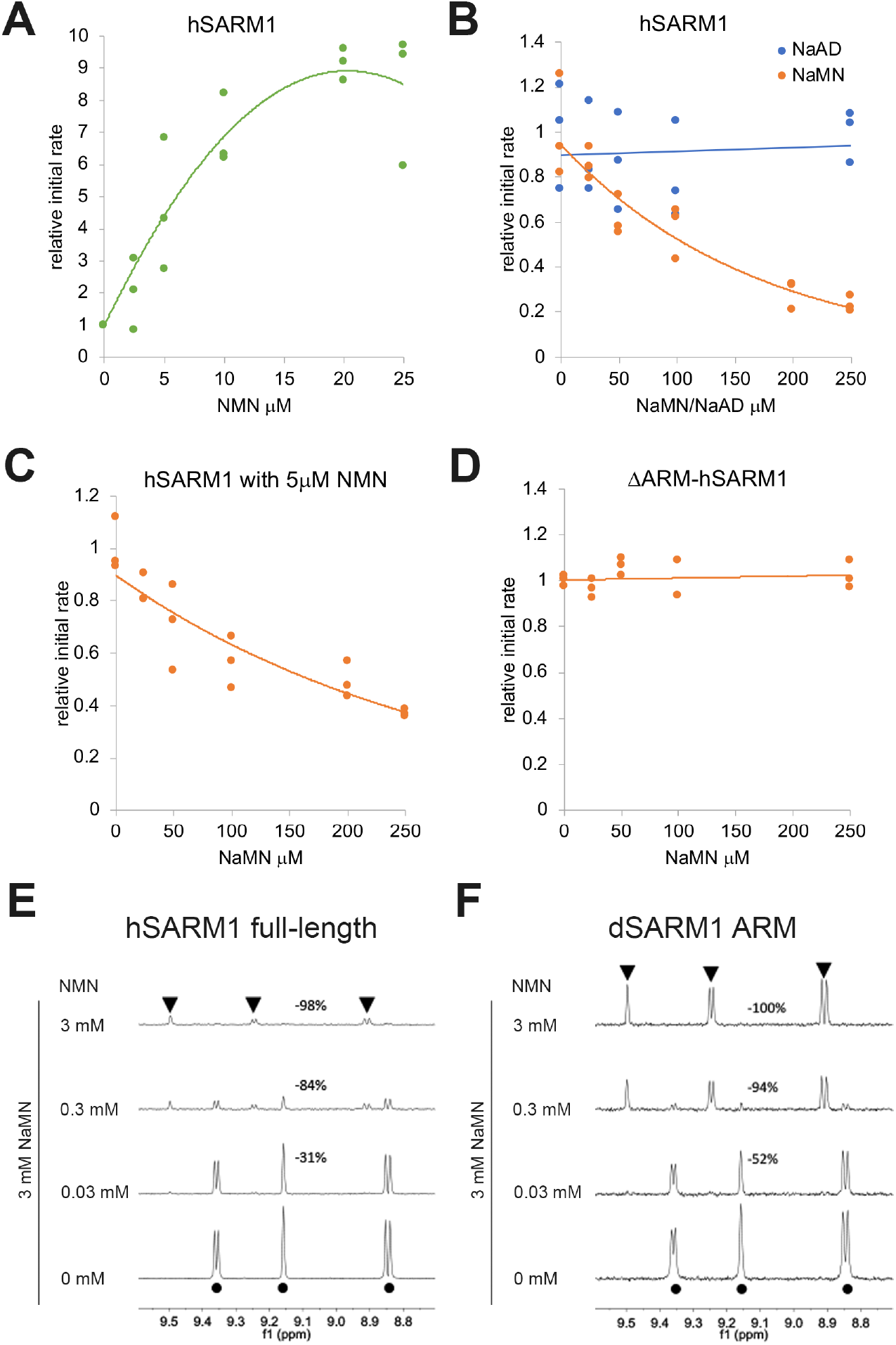
NaMN inhibits SARM1 NAD^+^ hydrolase activity. A-C) In-vitro SARM1 NAD^+^ hydrolase assay using full-length human SARM1 without the mitochondrial localization signal (hSARM1), using different concentrations of NMN or NaMN. Initial rates of the NAD^+^ hydrolase reaction normalized to control (0 μM NMN or NaMN) are shown. (A) NMN activated hSARM1 with Ka = 8.7 μM, (B) NaMN inhibited SARM1 with IC_50_ of 93.3 μM. NaAD did not show inhibition or activation of hSARM1 NAD^+^ hydrolase activity. (C) NaMN inhibited SARM1 with IC_50_ of 147.2 μM. (D) NaMN up to 250 μM did not show inhibition or activation of NAD^+^ hydrolase activity of hSARM1 lacking the N-terminal ARM domain (ΔARM-hSARM1). E-F) Expansions of saturation-transfer difference (STD) NMR spectra showing E) NaMN binding to 7.35 µM hSARM1 in the absence and presence of NMN and F) NaMN binding to 50 µM *Drosophila* SARM1 ARM domain (dSARM1 ARM) in the absence and presence of NMN. Four samples with 3 mM NaMN and increasing concentrations of NMN (from bottom to top: 0, 0.03 mM, 0.3 mM, and 3 mM) are shown. “•” indicates NaMN signals while “▾” indicates NMN signals. The percentage decrease of NaMN signals relative to no-NMN control is labelled.

Next, we directly assessed NaMN binding to SARM1 using saturation-transfer difference (STD) NMR(Figley et al., 2021). STD NMR assays demonstrated that NaMN binds to hSARM1and the *Drosophila* SARM1 ARM domain (dSARM1 ARM) (Figure 5E, F). Furthermore, NaMN binding to hSARM1 was decreased by 31% and 84% upon addition of NMN, at ratios of NMN:NaMN = 1:100 and 1:10, respectively. Similarly, NaMN binding to dSARM1 ARM was also inhibited by 52% and 94% at ratios of NMN:NaMN = 1:100 and 1:10 respectively. NMN completely abolished NaMN binding to hSARM1 and dSARM1 ARM when the two metabolites are present at equal concentrations, demonstrating that NaMN and NMN are binding to the same allosteric site (Figure 5E, F). The competition between NMN and NaMN, together with the lower affinity of NaMN for binding, explains why NMN levels must be reduced in order to observe NaMN-mediated (i.e. the addition of NaR) axonal protection.

To shed light on the mechanism of NaMN-mediated axonal protection, we determined the crystal structure of the NaMN-bound ARM domain of Drosophila SARM1 (dSARM1ARM) (PDB: 7RTC, Figure 6, Table 1). Previously, we determined the crystal structure of dSARM1ARM bound to the activating ligand NMN (Figley et al., 2021). The generation of the NaMN-bound ARM structure allowed us to analyze the essential structural differences induced by an inhibitory vs. an activating ligand. The NaMN-bound protein displays a right-handed superhelix, similar to ligand-free dSARM1 ARM (RMSD = 0.5 Å, PDB: 7LCY) (Figure 6A). The ligand NaMN binds to the allosteric site in the ARM domain and shares a similar binding mode with NMN (Figure 6B). NaMN is structurally very similar to NMN, differing only at the C3 position of the pyridine, where the amide group in NMN is replaced by the carboxyl group in NaMN. This small change induces a slightly more open conformation of the domain, with its C-terminus rotated away from the N-terminus by ∼11° (RMSD=1.4 Å) when compared to the NMN-bound dSARM1ARM (PDB: 7LCZ) (Figley et al., 2021). We reasoned that this open conformation is achieved through two related interactions. First, due to their negative charges, the carboxyl and phosphate groups of NaMN are repelled from each other, so that the nicotinic acid portion is pushed towards the N-terminus of the protein by ∼25 °, compared to the nicotinamide portion of NMN. This allows formation of a hydrogen bond with H392, which does not occur in the compact NMN-bound structure. Second, the carboxyl group of NaMN cannot interact with the loop connecting ARM6 and 7 (residues 594-602). In the NMN:dSARM1 ARM structure, this loop interacts with the amide group of NMN through two hydrogen bonds (with H599 and G600) and drags the C-terminus toward the N-terminus (Figley et al., 2021). Hence, NaMN keeps the structure in a more open conformation. Although both ligands bind in a very similar fashion, NaMN stabilizes a more extended conformation of the protein, more similar to the ligand-free structure. This altered conformation holds SARM1 in the inactive state and prevents it from promoting axon degeneration.

**Table 1.**
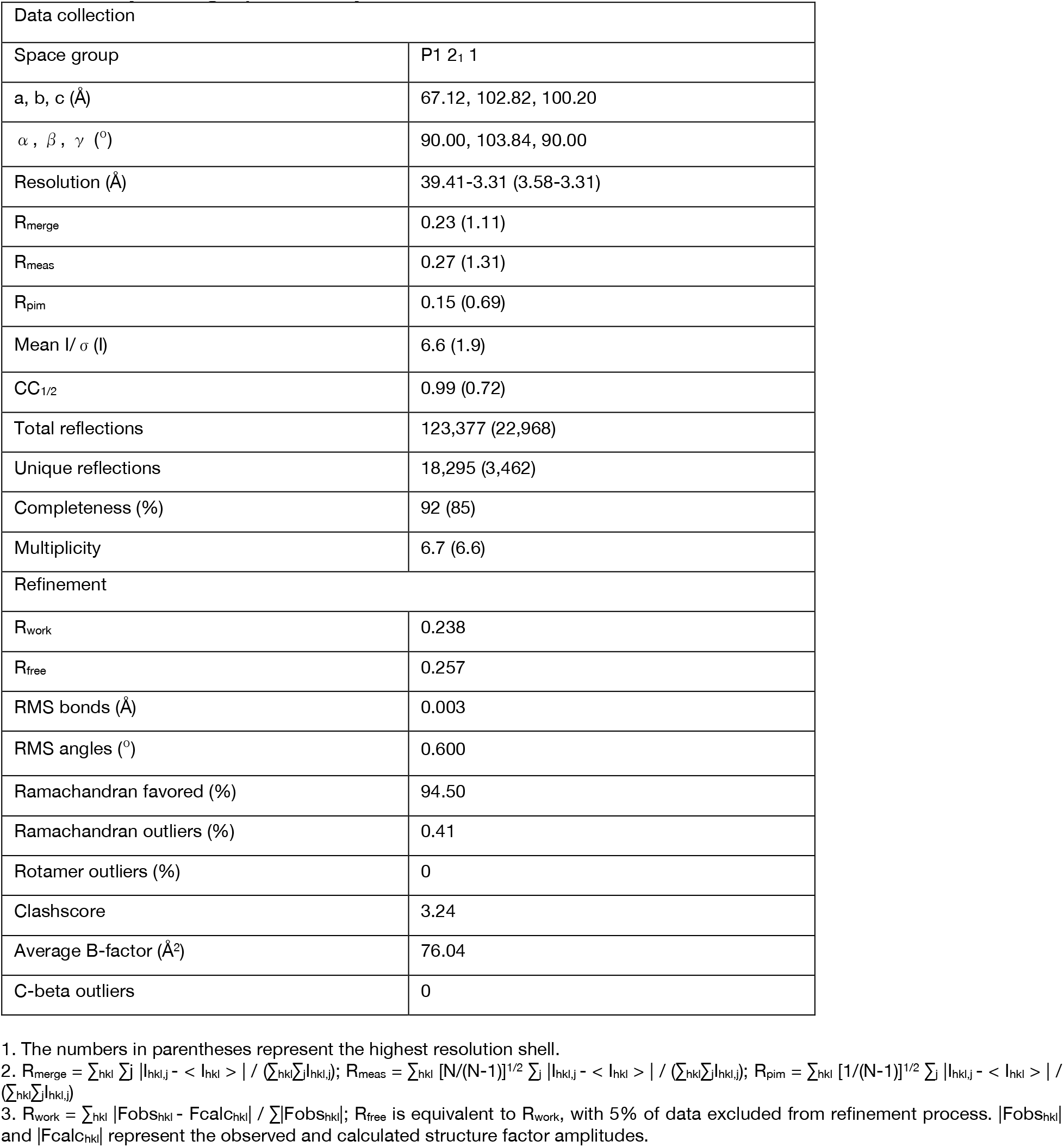
Crystallographic analysis.

**Figure 6.**
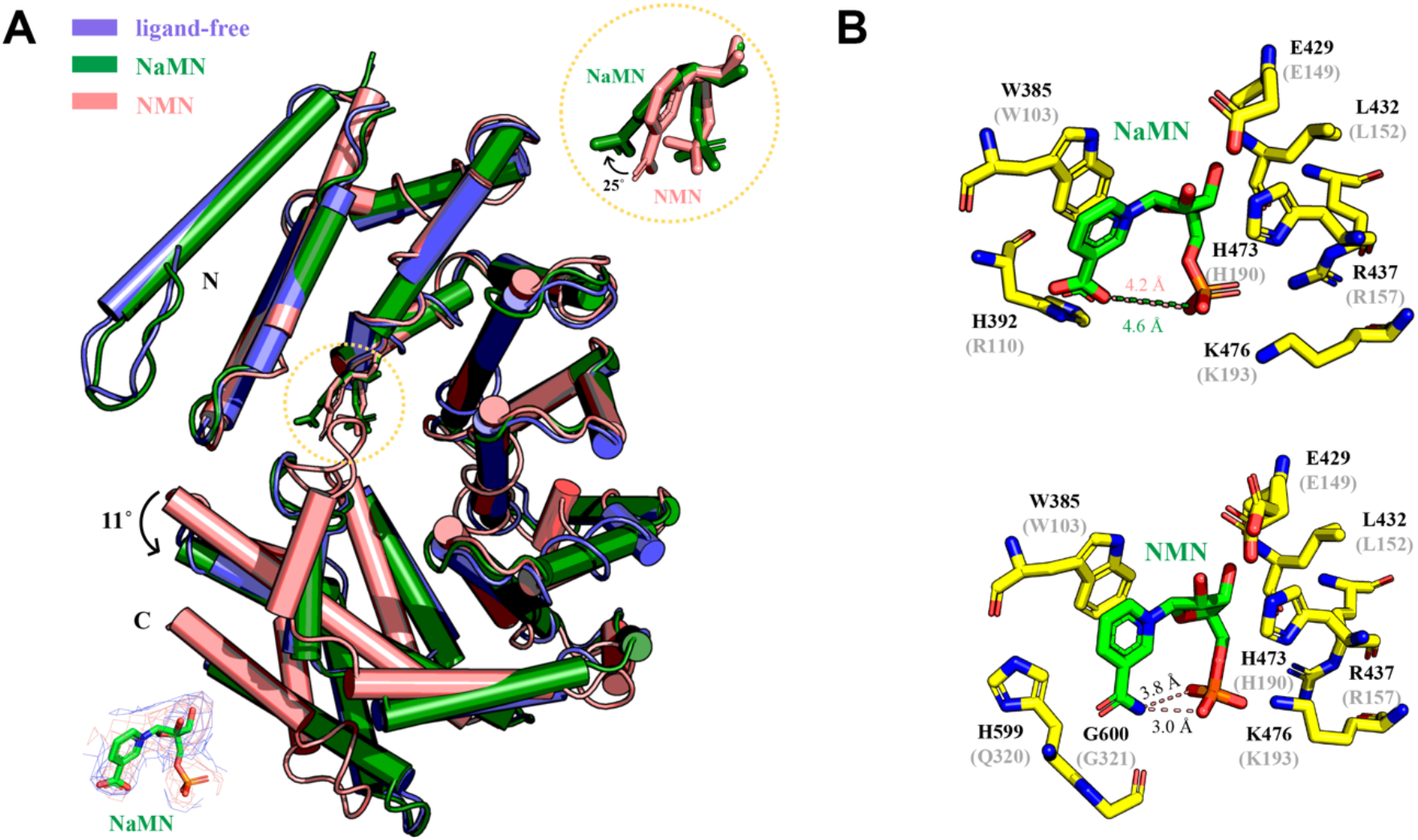
NaMN stabilizes an open conformation of dSARM1ARM. A) Superpositions of the N-terminal regions (residue 373-444) of NaMN-bound (green, PDB: 7RTC), NMN-bound (salmon, PDB: 7LCZ), and ligand-free dSARM1ARM (slate, PDB: 7LCY). Also shown are standard omit (deep-salmon mesh) and Polder (blue mesh) mFo-DFc maps for the NaMN molecule in dSARM1 ARM (bottom left), and zoom-in of the NaMN and NMN in stick representation (top right). B) Stick representation of the interaction between NaMN and dSARM1ARM (top), and between NMN and dSARM1ARM (bottom). The predicted NMN-binding residues in hSARM1 are shown in parentheses.

Having defined the structural basis for NaMN-mediated inhibition and discovered the molecular mechanism of axonal protection mediated by NaR plus FK866, we next addressed the mechanism of protection for the expression of *E. coli* NMN deamidase, which catalyzes the conversion of NMN to NaMN and potently blocks injury-induced axon degeneration both *in vitro* and *in vivo* (Di Stefano et al., 2017; 2015; Sasaki et al., 2016). Prior studies attributed NMN deamidase mediated protection exclusively to reduction in NMN levels; however, its axoprotective effects are dramatically greater than those observed with FK866 treatment, which also decreases NMN levels but does not increase NaMN levels. Hence, we hypothesized that NaMN also plays a role in long-term axonal protection mediated by NMN deamidase. If this were the case, then elevation of axonal NMN or reduction of NaMN should inhibit the long-term axonal protection by NMN deamidase just as it inhibits NaMN and FK866-mediated axonal protection. To boost NMN without the confound of a concomitant increase in NAD^+^ or NaMN 1) we applied NR, a cell permeable precursor of NMN, to axons 24 hours post-axotomy when the majority of NMNAT2 is lost and the conversion of NMN to NAD^+^ is suppressed (Figure 7A); and 2) we limited the amount of NMN deamidase in the axon to minimize the conversion of NMN to NaMN. To limit axonal NMN deamidase, protein transduction using lentivirus particles was used. We previously showed that the cytosolic form of NMNAT1 (cytNMNAT1) is able to block axon degeneration when it is introduced into axons as a protein using virus particles containing cytNMNAT1 (Sasaki and Milbrandt, 2010). To test if NMN deamidase also provides strong axonal protection via protein transduction, NMN deamidase virus particles were applied to the culture medium 10 minutes after axotomy (Figure 7A). We found that axonal degeneration was strongly inhibited for at least 48 hours (Figure 7B). Axonal metabolite measurements showed a significant elevation of NaMN at 24 hours post-axotomy in NMN deamidase protein-transduced axons compared with control or FK866 treated axons (Figure 7C, D). In contrast, the level of NMN was comparable to the control axons at 24 hours post axotomy and higher than FK866-treated axons at 0 and 24 hours post-axotomy. In addition, the NAD^+^ levels were not restored in NMN deamidase transduced axons compared with control axons at 0 or 24 hours post-axotomy. Hence, protein transduction of NMN deamidase provides much longer lasting protection than does FK866 (compare Figure 7B to Figure 1D), yet NMN levels are higher and NAD^+^ levels are comparable. Therefore, the protection afforded by NMN deamidase is not solely due to changes in the levels of NMN or NAD^+^.

**Figure 7.**
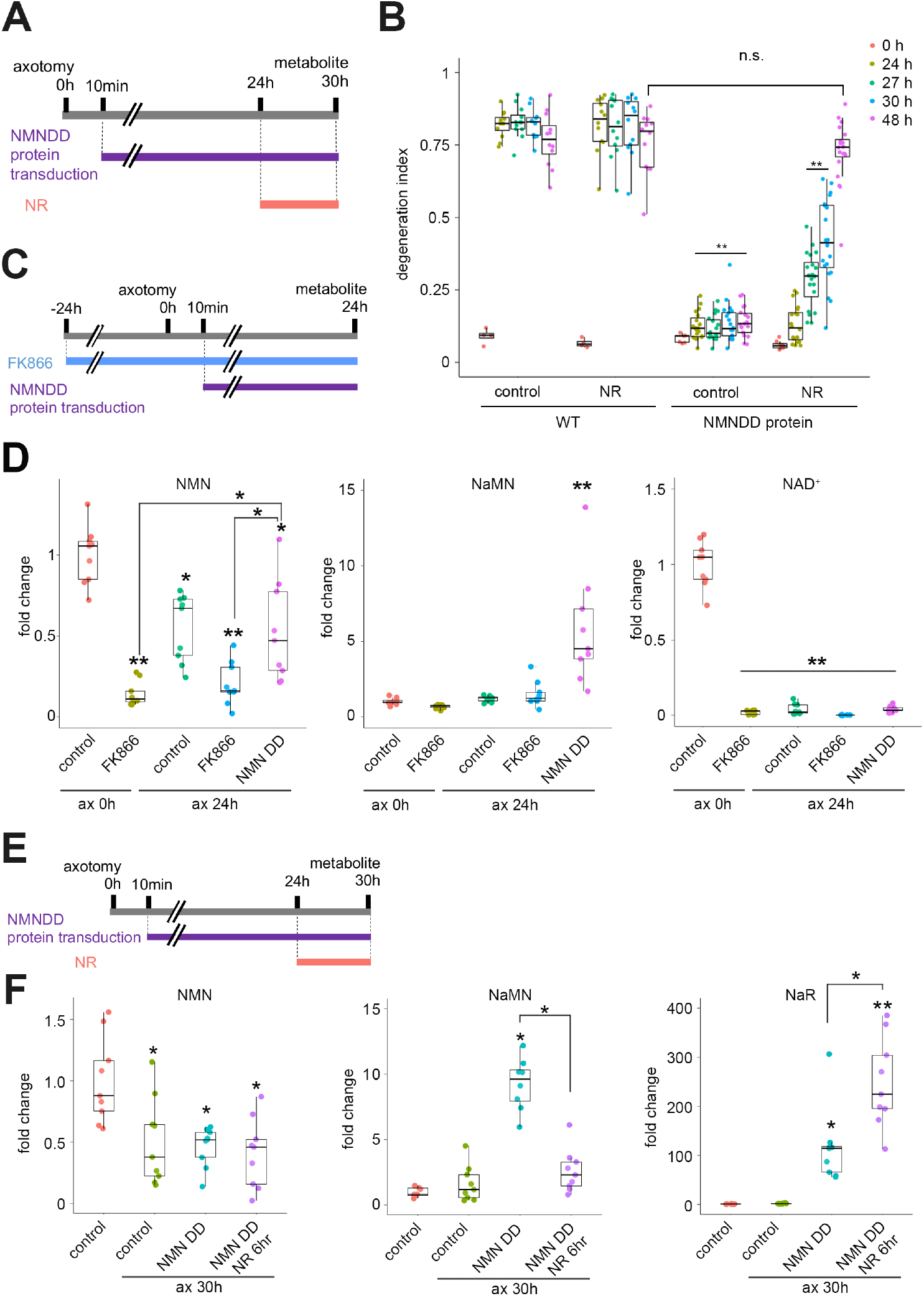
NMN deamidase supports prolonged axonal protection by decreasing NMN and increasing NaMN. A) Time scheme showing the timing of axotomy, chemical addition (NaR 100 μM), and axon degeneration assay used in (B). B) Axonal degeneration of wild-type neurons from 0 to 48 h post-axotomy with treatments described in (A) was quantified. NR (100 μM) addition at 24 hours post-axotomy to the NMN deamidase protein transduced axons showed axon degeneration by 48 hours post-axotomy. Statistical analysis was performed by two-way ANOVA with a Tukey post-hoc test. F(3, 256)=946.7, p<2.0×10^−16^ among groups. **p<1×10^−5^ denotes the significant difference compared with the degeneration index of control at the same time post-axotomy. N.S. denotes no significant difference. C) Time scheme showing the timing of axotomy, addition of FK866 (100 nM) or NMN deamidase (NMNDD) protein transduction and metabolite measurement. D) Axonal metabolites 0 or 24 h post-axotomy from neurons treated with FK866 or NMNDD protein transduction. NMN levels in the presence of NMNDD protein are not lower than FK866-treated axons with or without axotomy. Statistical analysis was performed by one-way ANOVA with Tukey post-hoc test. F(4, 40)=27.1, p=6.4×10^−11^ for NMN, F(4, 40)=84, p=13.8×10^−7^ for NaMN, F(4, 40)=340, p<2.0×10^−16^ for NAD^+^ among groups. **p<1 × 10^−5^ and *p<0.05 denote significant difference compared with wild-type control axons. E) Time scheme showing the timing of axotomy, addition of NR (100 μM), NMN deamidase (NMNDD) protein transduction, and metabolite measurement. F) Axonal metabolites 0 or 30 h post-axotomy with or without NMNDD protein transduction at 24 h post-axotomy and plus/minus NR (100 μM). NaMN was reduced without changing NMN after NR addition to NMNDD protein-transduced axons. Statistical analysis was performed by one-way ANOVA with a Tukey post-hoc test. F(3, 32)=7.6, p= 5.4 ×10^−4^ for NMN, F(2, 32)=6.6 , p=1.2×10^−3^ for NaMN , F(3, 32)=35.1, p<3.1×10^−10^ for NAD^+^ among groups. **p<1 × 10^−5^ and *p<0.05 denote significant differences compared with wild-type control axons.

Next, we tested whether NR treatment to injured axons would reverse NMN deamidase-mediated protection, similar to what we observed in axons preserved by combined NaR/FK866 treatment. We found that NR (100 μM) addition 24 hours post-axotomy potently stimulated axon degeneration in axons transduced with NMN deamidase protein (Figure 7B). We then measured axonal metabolites in NMN deamidase protein transduced axons in the presence of NR. NMN deamidase containing virus particles were added 10 minutes post-axotomy, NR was added 24 hours post-axotomy, and axonal metabolites were measured 6 hours later (30 hours post-axotomy, Figure 7E). Interestingly, axonal NMN was not increased, demonstrating that the NMN deamidase can still suppress NMN levels even following the addition of NR. However, NR addition did lead to a large decrease in NaMN to the levels of control axons lacking NMN deamidase, and also induced a robust increase in the levels of NaR (Figure 7F). While this result was initially surprising, it is likely explained by the generation of NaMN by NMN deamidase, followed by the dephosphorylation of NaMN to NaR by the purine nucleoside phosphorylase family of phosphatases (Belenky et al., 2009). We speculate that NR supplementation competes with NaR for access to the NRK enzymes that convert NaR to NaMN, leading to unopposed phosphatase activity that keeps NaMN low and NaR high. As the addition of NR blocks axon protection mediated by protein transduction of NMN deamidase, these findings strongly suggest that NMN deamidase inhibits axon degeneration via two steps: by reducing levels of the SARM1 activator, NMN, and by elevating levels of the SARM1 inhibitor, NaMN. These steps are also carried out by a combination of FK866 (NMN reduction) and NaR (NaMN increase) to inhibit axon degeneration. These findings revise our understanding of NMN deamidase and FK866/NaR mediated axon protection and highlight the potential of NMN analogs as key modulators of SARM1 activation.

## Discussion

The mechanism of SARM1 activation was recently elucidated, identifying SARM1 as a metabolic sensor regulated by the ratio of NMN-to-NAD^+^ via their competitive binding to an allosteric pocket in the SARM1 autoinhibitory domain (Figley et al., 2021). Here we demonstrate that NaMN, the deamidated form of NMN, competes with NMN for binding to the SARM1 allosteric pocket and thereby inhibits SARM1 activation. The identification of NaMN as a SARM1 inhibitor provides a new mechanistic understanding for previously described treatments that provide strong protection of of injured axons (Di Stefano et al., 2017; 2015; Liu et al., 2018). Notably, these treatments have a dual mechanism of action: they both decrease levels of the activator NMN and simultaneously produce NaMN, which inhibits SARM1 by competing with NMN at the SARM1 allosteric site and stabilizing the extended, inactive conformation. The identification of a third NAD^+^ metabolite binding and regulating SARM1 underscores the central role of NAD^+^ metabolism in regulating the activation of SARM1 NAD^+^ hydrolysis and subsequent neurodegeneration. Moreover, these findings highlight the potential of SARM1 allosteric inhibitors as novel therapeutics for the treatment of neurodegeneration.

Here we identify NaMN as a SARM1 inhibitor and reveal the mechanism of strong axonal protection provided by the combination of NaR and FK866 or the neuronal expression of NMN deamidase. A series of elegant experiments from the Coleman, Conforti, and Zhao labs highlighted the role of NMN in promoting SARM1-dependent axon degeneration (Di Stefano et al., 2017; 2015; Gilley and Coleman, 2010; Zhao et al., 2019), and our recent demonstration that NMN binds to an allosteric pocket and activates SARM1 provides the mechanistic explanation for these observations (Figley et al., 2021). However, a few inconsistencies persisted in the literature. Most strikingly, two mechanisms that each reduce NMN levels yield dramatically different degrees of axonal protection (Di Stefano et al., 2015; Sasaki et al., 2009). Inhibition of NMN synthesis by treatment with the NAMPT inhibitor FK866 results in less than one day of protection for severed axons, while the conversion of NMN to NaMN by the neuronal expression of the bacterial enzyme NMN deamidase provides at least 72 hours of protection (Di Stefano et al., 2015; Sasaki et al., 2016). While the simplest explanation for this discrepancy would be that the NMN deamidase lowers NMN levels more than does FK866, we show here that FK866 can lower NMN levels even more than the NMN deamidase while providing much poorer protection. The second surprising finding was the dramatic enhancement of FK866-mediated protection by the addition of NaR in axons injured by the neurotoxin vincristine (Liu et al., 2018). While it was originally suggested that NaR could be bypassing the NAD^+^ salvage pathway to generate NAD^+^ in these injured axons, we questioned this explanation because the generation of axonal NAD^+^ requires the NAD^+^ biosynthetic enzyme NMNAT2, which is rapidly lost in injured axons following both axotomy and vincristine treatment (Gilley and Coleman, 2010; Summers et al., 2019). Here we resolve these discrepancies by demonstrating that: 1) both of these strongly protective treatments lead to the generation of NaMN; 2) NaMN directly binds to SARM1 and promotes the open ARM domain conformation found in inactive SARM1 3) NaMN inhibits SARM1 activation; 4) NaMN competes with NMN for SARM1 binding; and 5) NaMN binds SARM1 more weakly than does NMN, explaining why NMN levels must be lowered to observe protection by NaMN.

We only observe NaMN-mediated protection when NMN levels are artificially lowered, raising the question of if and when this mechanism might be relevant to axonal maintenance in an untreated neuron. NMN is generated by the NAD^+^ salvage pathway, while NaMN is generated by both the *de novo* and Preiss-Handler NAD^+^ synthesis pathways. In mammals, the salvage pathway is generally predominant, and we certainly find that to be true in the DRG neurons tested here. However, there may be neuronal cell types or conditions in which there is a relative preference for the *de novo* or Preiss-Handler pathways, which would boost the NaMN/NMN ratio and thereby decrease the likelihood of SARM1 activation. For example, in a neuronal subtype with abundant NaPRT using the common vitamin B3 analog nicotinic acid would lead to high levels of NaMN. We speculate that differences in NAD^+^ biosynthetic routes are a candidate mechanism for differential vulnerability of neurons due to the antagonistic effects of NMN and NaMN on SARM1 activation.

The SARM1 allosteric regulatory pocket binds NaMN, NAD^+^, and NMN, with NaMN and NAD^+^ inhibiting NMN binding and thereby antagonizing SARM1 activation (Figley et al., 2021; Jiang et al., 2020; Sporny et al., 2020). Hence, the SARM1 allosteric pocket is relatively promiscuous, binding diverse components of the NAD^+^ metabolome. This suggests that additional endogenous and exogenous molecules could bind this pocket and regulate SARM1 activation. Indeed, multiple neurotoxins binding the SARM1 allosteric pocket have recently been identified(Loreto et al., 2021; Wu et al., 2021). This relative promiscuity gives hope that small molecules could be developed that would bind the pocket much more tightly than either NaMN or NAD^+^ to block NMN binding and to inhibit SARM1 activation. The ability to target both SARM1 activation at the allosteric site and SARM1 activity within the TIR enzymatic pocket may enable powerful synergistic approaches to inhibit SARM1 for the prevention and treatment of neurodegenerative diseases (Bosanac et al., 2021; Hughes et al., 2021; Li et al., 2021).

## Materials and Methods

Chemicals: FK866 was obtained from National Institute of Mental Health Chemical Synthesis and Drug Supply Program (MH number F-901). Nicotinic acid riboside was purchased from Santa Cruz Biotechnology (sc-478242). Nicotinamide riboside (NR) was a gift from ChromaDex, Inc.. NAD^+^ (Sigma, N1636), NMN (Sigma, N3501), cADPR (Sigma, C7344) were purchased from Sigma.

Mouse studies: CD1 mice (gestation day 12.5; Charles River Laboratories) were housed (12 h dark/light cycle and less than 5 mice per cage) and used under the direction of institutional animal study guidelines at Washington University in St. Louis. NMRK1 knockout mice: Nmrk1^tm1a(KOMP)Wtsi^ (NRK1^-/-^) and NMRK2 knockout mice: Nmrk2 ^tm1.1(KOMP)Vlcg/J^ (NRK2^-/-^) were obtained from Wellcome Trust Sanger Institute and Jackson laboratory respectively and were used to generated NRK1^-/-^:NRK2^-/-^ (NRK1/2 dKO, Supplemental Figure 1).

DRG neuronal culture and axon degeneration assay: mouse DRG neuron culture was performed as described before (Sasaki et al., 2016). Mouse dorsal root ganglion (DRG) was dissected from embryonic days 13.5 CD1 mouse embryo (50 ganglia per embryo) and incubated with 0.05% trypsin solution containing 0.02% EDTA (Gibco) at 37°C for 15 min. Then cell suspensions were triturated by gentle pipetting and washed 3 times with the DRG growth medium (Neurobasal medium (Gibco) containing 2% B27 (Invitrogen), 100 ng/ml 2.5S NGF (Harlan Bioproduts), 1 µM uridine (Sigma), 1 µM 5-fluoro-2’-deoxyuridine (Sigma), penicillin, and streptomycin). Cells were suspended in DRG growth medium at a ratio of 50 µl medium/50 DRGs. The cell density of these suspensions was ∼1×10^7^ cells/ml. Cell suspensions (1 µl/96 well, 5 µl/24 well) were placed in the center of the well using either 96- or 24-well tissue culture plates (Corning) coated with poly-D-Lysine (0.1 mg/ml; Sigma) and laminin (3 µg/ml; Invitrogen). Cells were allowed to adhere in humidified tissue culture incubator (5% CO_2_) for 15 min and then DRG growth medium was gently added (100 µl/96 well, 500 µl/24 well). Lentivirus transfer vector constructs harboring cDNAs including Venus, cytNMNAT1 (mouse), NRK1 (mouse), and NMN deamidase (E. coli) were previously described (Sasaki et al., 2016). Lentiviruses were added (1-10×10^3^ pfu) at 1–2 days *in vitro* (DIV) and metabolites were extracted or axon degeneration assays (Sasaki et al., 2009) were performed at 6–7 DIV. Data were derived from minimum three biological replicates using independent batch of DRG cells originated from independent pregnant mice and minimum three technical replicates were included in each biological replicate. For protein transduction of NNM deamidase, lentivirus particles (5-50×10^3^) were added at indicated time post axotomy. When using 24 well DRG cultures, 50% of the medium was exchanged for a fresh medium at DIV4. For axotomy experiments, cell bodies and axons were separated by a microsurgical blade.

Metabolite extraction: At DIV6, tissue culture plates were placed on ice and culture medium replaced with ice-cold saline (0.9% NaCl in water, 500 µl per well). For collection of axonal metabolites, cell bodies were removed using a pipette. For collection of intracellular metabolites, saline was removed and replaced with 160 µL ice cold 50% MeOH in water. The axons were incubated for a minimum of 5 min on ice with the 50% MeOH solution and then the solution was transferred to tubes containing 50 µl chloroform on ice, shaken vigorously, and centrifuged at 20,000 x g for 15 min at 4°C. The clear aqueous phase (140 µl) was transferred into a microfuge tube and lyophilized under vacuum. Lyophilized samples were stored at -20°C until measurement.

Metabolite measurements: Lyophilized samples were reconstituted with 5 mM ammonium formate (15 µl), centrifuged (13,000 x g, 10 min, 4°C), and 10 µl clear supernatant was analyzed. NMN, NAD^+^, and cADPR were measured using LC-MS/MS (Hikosaka et al., 2014). Samples were injected into a C18 reverse phase column (Atlantis T3, 2.1 × 150 mm, 3 µm; Waters) using HPLC (Agilent 1290 LC) at a flow rate of 0.15 ml/min with 5 mM ammonium formate for mobile phase A and 100% methanol for mobile phase B. Metabolites were eluted with gradients of 0–10 min, 0–70% B; 10– 15 min, 70% B; 16–20 min, 0% B. The metabolites were detected with a triple quad mass spectrometer (Agilent 6470 MassHunter; Agilent) under positive ESI multiple reaction monitoring (MRM) using parameters specific for each compound (NAD^+^, 664>428, fragmentation (F) = 160 V, collision (C) = 22 V, and cell acceleration (CA) = 7 V: NMN 335 > 123, F=135 V, C=8 V, CA = 7 V: cADPR, 542>428, F = 100 V, C = 20 V, and CA = 3 V: NaMN 336>124, F=80 V, C=10V, CA = 7 V: NaAD 669>428, F=150 V, C=20 V, CA = 7 V: NaR, 256>124, F=35 V, C=10V, CA = 3 V: NR 255 >123, F=130 V, C=20 V, CA = 1 V). Serial dilutions of standards for each metabolite in 5 mM ammonium formate were used for calibration. Metabolites were quantified by MassHunter quantitative analysis tool (Agilent) with standard curves and normalized by the protein. Data were derived from a minimum of three biological replicates using independent batches of DRG cells dissected from independent pregnant mice and a minimum of three technical replicates were included in each biological replicate.

Western blot: DRG cell suspension (1×10^7^ cells/ml, 5μl) was spotted in the center of 24-well and cultured for 6 to 7 days as described above. Cell bodies were separated from axons using the micro surgical blade and were removed from the culture vessel by aspirating with P200 pipet. After indicated time post axotomy, culture medium was removed, and axons were collected with 300 μL of 0.9% NaCl in water and transferred to a 1.5 ml microcentrifuge tube. After centrifugation (12,000xg, 5min), the supernatant was removed and added 100 μl 10mM phosphate buffer (pH 7.4) containing 138mM NaCl, 2.7mM KCl, and protease inhibitor cocktail (cOmplete, Roche) and axons were lysed by sonication and protein concentration was measured using BCA Protein Assay Kit (Pierc). Then axon lysates were mixed with sample buffer (375mM Tris-HCl (pH6.8), 9% SDS, 50% glycerol, and 0.03% bromophenol blue) and about 10 μg of the lysates were used for western blotting. SDS-PAGE was performed using Bolt™ 4 to 12%, Bis-Tris (Invitrogen) with Bolt MOPS SDS Running Buffer (Invitrogen) and proteins were transferred to nitrocellulose membrane (Amersham, 10600002). Membranes were incubated with blocking solution (5% BSA in PBS containing 0.1% Triton X-100 (PBS-T)) for one hour then with NMNAT2 antibody (Santa Cruz, sc-515206, 1:1000 dilution in blocking solution) overnight at 4°C. Membranes were washed in PBS-T and incubated with HRP-conjugated secondary antibody (Jackson Immuno Research, 115-007-003, 1:5000 dilution in PBS-T) for one hour and developed with chemiluminescent reaction (VWR, 490005-008).

*In-vitro* SARM1 NAD^+^ hydrolase assay: human SARM1 without mitochondria targeting sequence and with N-terminal tandem StrepTag II was expressed in HEK293T cells expressing NRK1 supplemented with NR (Essuman et al., 2017) and purified with MagStrep (Strep-Tactin) type 3 XT beads suspension (IBA Lifesciences). The purified protein attached on beads was incubated with NAD^+^ (10 μM for Figure 5 B, C and 50 μM for Figure 5A) for 0, 5, 15, and 30 minutes at 25°C in the presence or absence of various concentrations of NMN, NaMN, or NMN plus NaMN that were preincubated with SARM1 before NAD^+^ addition. Metabolites were extracted with the chloroform/methanol method, lyophilized, and measured by HPLC as previously described (Essuman et al., 2017).

Saturation-transfer difference (STD) NMR: both hSARM1 and dSARM1 ARM were produced as per Figley et al (Figley et al., 2021), and STD NMR was performed in a similar fashion as described previously (Figley et al., 2021). Briefly, samples for STD NMR were prepared in a total volume of 200 μl, with each sample consisting of 175 μl HBS buffer, 20 μL D_2_O, and 5 μl DMSO-d6. STD NMR spectra were acquired with a Bruker Avance 600 MHz NMR spectrometer equipped with ^1^H/^13^C/^15^N triple resonance cryoprobe at 298 K. The pulse sequence STDDIFFGP19.3, built-in within the TopSpin™ program (Bruker), was employed to acquire STD NMR spectra (Mayer and Meyer, 1999). This pulse sequence consists of a 3-9-19 water suppression pulse, the parameters of which were obtained from the water suppression pulse program (P3919GP), to suppress the resonance from H_2_O. The on-resonance irradiation was set close to protein resonances at 0.8 ppm, whereas the off-resonance irradiation was set far away from any protein or ligand resonances at 300 ppm. A relaxation delay of 4 s was used, out of which a saturation time of 3 s was used to irradiate the protein with a train of 50 ms Gaussian shaped pulses. The number of scans was 512. All spectra were processed by TopSpin (Bruker) and Mnova 11 (Mestrelab Research).

Crystallization and crystal structure determination: Crystallization of dSARM1 ARM (residues 315-678) was performed as previously described (Figley et al., 2021). The crystals were soaked in 1.7 M sodium malonate (pH 5.8) with 20 mM NaMN (Sigma, N7764) at 18 °C overnight, protected with Paratone-N and flash-cooled in liquid nitrogen. X-ray diffraction data were collected using the wavelength of 0.95373 Å at the Australian Synchrotron MX2 beamline (Aragão et al., 2018) and processed with the program autoPROC (Vonrhein et al., 2011). The structure (PDB: 7RTC) was determined using the molecular replacement approach, with the ligand-free dSARM1ARM (PDB: 7LCY) structure as a search model using Phaser (McCoy et al., 2007), and refined and manually (re)-built using phenix.refine (Afonine et al., 2012) and Coot (Emsley and Cowtan, 2004), respectively.

## Acknowledgements

The work was supported by the National Health and Medical Research Council (NHMRC grants 1107804 and 1160570 to B.K. and T.V., 1071659 to B.K., and 1196590 to T.V.), Needleman Center for Neurometabolism and Axonal Therapeutics (J.M. and A.D.), the Australian Research Council (ARC) Laureate Fellowship (FL180100109 to B.K.), and Future Fellowship (FT200100572 to T.V.), and National Institutes of Health grants (R01CA219866 and RO1NS087632 to A.D. and J.M. and RF1AG013730 to J.M.). We thank Matthew Figley for sharing unpublished observations. We thank Kelli Simburger and Tim Fahrner for support with molecular cloning, Cassidy Menendez, Rachel McClarney and Mihir Vohra for animal husbandry and maintenance, Alicia Neiner for processing LC-MS-MS samples, and Tamim Mosaiab and Veronika Masic for support with NMR experiments and protein production. We thank the use of the University of Queensland Remote Operation Crystallization and X-ray (UQROCX) Facility at the Centre for Microscopy and Microanalysis, and Australian Synchrotron MX facility. We thank members of the Kobe, Ve, DiAntonio and Milbrandt labs for helpful discussions.

## Author Contributions

Conceptualization, Y.Sasaki, A.D., J.M. ;Investigation, Y.Sasaki., J.Z., Y.S., W.G., B.K., T.V., A.D., J.M.; Writing –Original Draft, Y.Sasaki, A.D., J.M. ; Writing – Review & Editing, All authors; Funding Acquisition, B.K., T.V., A.D., J.M.; Supervision, B.K., T.V., A.D., J.M.

---

**Supplemental Figure 1.**
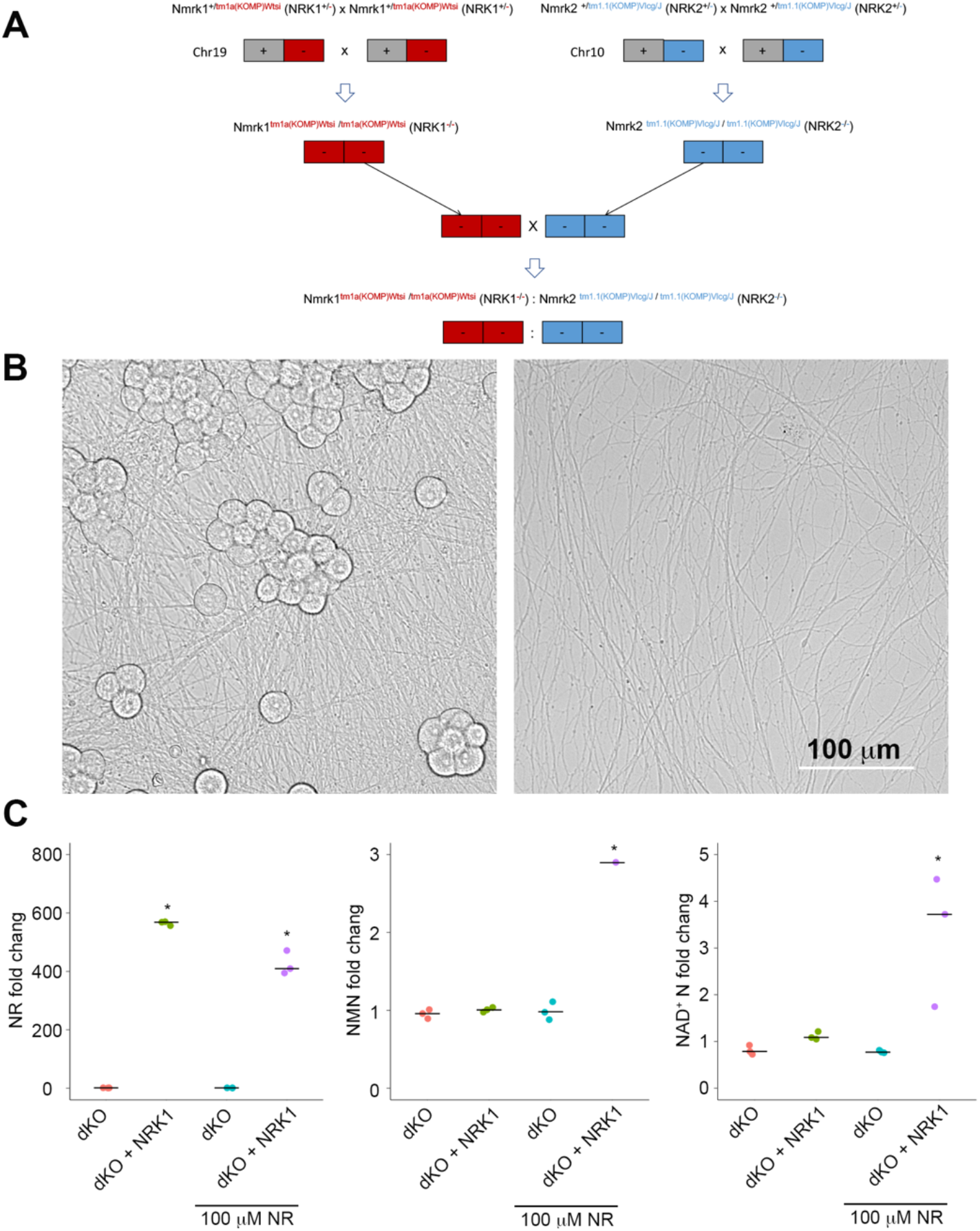
NRK1/2 dKO mice and neurons are viable. A) A breeding scheme showing the generation of Nmrk1:Nmrk2 double knockout (NRK1/2 dKO) mice. B) Images showing DIV7 DRG neuronal cell bodies and axons from NRK1/2 dKO mice. Scale: 100 μm. C) Metabolite analysis from NRK1/2 dKO neurons expressing Venus (dKO) or mouse NRK1 (dKO+NRK1) via lentivirus. NMN or NAD^+^ were low in NRK1/2 dKO neurons with or without NR addition (100 μM, 24 h). Exogenously expressed NRK1 produces high NMN and NAD^+^ in NRK1/2 dKO neurons. Statistical analysis was performed by one-way ANOVA with a Tukey post-hoc test. F(3, 8)=31.9, p=8.5×10^−5^ for NR, F(3, 8)=592.4, p=9.9×10^−10^ for NMN, and F(3 ,44)=8.8 P=6.4 ×10^−3^ for NAD^+^. *p<0.05 denotes the significant difference compared with metabolites of the NRK1/2 dKO control neurons.

## References

Afonine, P.V., Grosse-Kunstleve, R.W., Echols, N., Headd, J.J., Moriarty, N.W., Mustyakimov, M., Terwilliger, T.C., Urzhumtsev, A., Zwart, P.H., Adams, P.D., 2012. Towards automated crystallographic structure refinement with phenix.refine. Acta Crystallogr D Biol Crystallogr 68, 352–367. doi:10.1107/S0907444912001308

Aragão, D., Aishima, J., Cherukuvada, H., Clarken, R., Clift, M., Cowieson, N.P., Ericsson, D.J., Gee, C.L., Macedo, S., Mudie, N., Panjikar, S., Price, J.R., Riboldi-Tunnicliffe, A., Rostan, R., Williamson, R., Caradoc-Davies, T.T., 2018. MX2: a high-flux undulator microfocus beamline serving both the chemical and macromolecular crystallography communities at the Australian Synchrotron. J Synchrotron Radiat 25, 885–891. doi:10.1107/S1600577518003120

Belenky, P., Christensen, K.C., Gazzaniga, F., Pletnev, A.A., Brenner, C., 2009. Nicotinamide riboside and nicotinic acid riboside salvage in fungi and mammals. Quantitative basis for Urh1 and purine nucleoside phosphorylase function in NAD+ metabolism. J. Biol. Chem. 284, 158–164. doi:10.1074/jbc.M807976200

Bloom, A.J., Mao, X., Strickland, A., Sasaki, Y., Milbrandt, J., DiAntonio, A., 2021. Constitutively active SARM1 variants found in ALS patients induce neuropathy. bioRxiv 2021.04.16.439886. doi:10.1101/2021.04.16.439886

Bosanac, T., Hughes, R.O., Engber, T., Devraj, R., Brearley, A., Danker, K., Young, K., Kopatz, J., Hermann, M., Berthemy, A., Boyce, S., Bentley, J., Krauss, R., 2021. Pharmacological SARM1 inhibition protects axon structure and function in paclitaxel-induced peripheral neuropathy. Brain. doi:10.1093/brain/awab184

Cetinkaya-Fisgin, A., Luan, X., Reed, N., Jeong, Y.E., Oh, B.C., Höke, A., 2020. Cisplatin induced neurotoxicity is mediated by Sarm1 and calpain activation. Scientific Reports 2016 6 10, 21889–12. doi:10.1038/s41598-020-78896-w

Di Stefano, M., Loreto, A., Orsomando, G., Mori, V., Zamporlini, F., Hulse, R.P., Webster, J., Donaldson, L.F., Gering, M., Raffaelli, N., Coleman, M.P., Gilley, J., Conforti, L., 2017. NMN Deamidase Delays Wallerian Degeneration and Rescues Axonal Defects Caused by NMNAT2 Deficiency In Vivo. Curr. Biol. 27, 784–794. doi:10.1016/j.cub.2017.01.070

Di Stefano, M., Nascimento-Ferreira, I., Orsomando, G., Mori, V., Gilley, J., Brown, R., Janeckova, L., Vargas, M.E., Worrell, L.A., Loreto, A., Tickle, J., Patrick, J., Webster, J.R.M., Marangoni, M., Carpi, F.M., Pucciarelli, S., Rossi, F., Meng, W., Sagasti, A., Ribchester, R.R., Magni, G., Coleman, M.P., Conforti, L., 2015. A rise in NAD precursor nicotinamide mononucleotide (NMN) after injury promotes axon degeneration. Cell Death and Differentiation 2014 22:5 22, 731–742. doi:10.1038/cdd.2014.164

Emsley, P., Cowtan, K., 2004. Coot: model-building tools for molecular graphics. Acta Crystallogr D Biol Crystallogr 60, 2126–2132. doi:10.1107/S0907444904019158

Essuman, K., Summers, D.W., Sasaki, Y., Mao, X., DiAntonio, A., Milbrandt, J., 2017. The SARM1 Toll/Interleukin-1 Receptor Domain Possesses Intrinsic NAD+Cleavage Activity that Promotes Pathological Axonal Degeneration. Neuron 93, 1334– 1343.e5. doi:10.1016/j.neuron.2017.02.022

Essuman, K., Summers, D.W., Sasaki, Y., Mao, X., Yim, A.K.Y., DiAntonio, A., Milbrandt, J., 2018. TIR Domain Proteins Are an Ancient Family of NAD+-Consuming Enzymes. Curr. Biol. 28, 421–430.e4. doi:10.1016/j.cub.2017.12.024

Figley, M.D., DiAntonio, A., 2020. The SARM1 axon degeneration pathway: control of the NAD+ metabolome regulates axon survival in health and disease. Current Opinion in Neurobiology 63, 59–66. doi:10.1016/j.conb.2020.02.012

Figley, M.D., Gu, W., Nanson, J.D., Shi, Y., Sasaki, Y., Cunnea, K., Malde, A.K., Jia, X., Luo, Z., Saikot, F.K., Mosaiab, T., Masic, V., Holt, S., Hartley-Tassell, L., McGuinness, H.Y., Manik, M.K., Bosanac, T., Landsberg, M.J., Kerry, P.S., Mobli, M., Hughes, R.O., Milbrandt, J., Kobe, B., DiAntonio, A., Ve, T., 2021. SARM1 is a metabolic sensor activated by an increased NMN/NAD+ ratio to trigger axon degeneration. Neuron 109, 1118–1136.e11. doi:10.1016/j.neuron.2021.02.009

Geisler, S., Doan, R.A., Cheng, G.C., Cetinkaya-Fisgin, A., Huang, S.X., Höke, A., Milbrandt, J., DiAntonio, A., 2019. Vincristine and bortezomib use distinct upstream mechanisms to activate a common SARM1-dependent axon degeneration program. JCI Insight 4, 135. doi:10.1172/jci.insight.129920

Geisler, S., Doan, R.A., Strickland, A., Huang, X., Milbrandt, J., DiAntonio, A., 2016. Prevention of vincristine-induced peripheral neuropathy by genetic deletion of SARM1 in mice. Brain 139, 3092–3108. doi:10.1093/brain/aww251

Gerdts, J., Brace, E.J., Sasaki, Y., DiAntonio, A., Milbrandt, J., 2015. SARM1 activation triggers axon degeneration locally via NAD+destruction. Science 348, 453–457. doi:10.1126/science.1258366

Gerdts, J., Summers, D.W., Sasaki, Y., DiAntonio, A., Milbrandt, J., 2013. Sarm1-mediated axon degeneration requires both SAM and TIR interactions. J. Neurosci. 33, 13569–13580. doi:10.1523/JNEUROSCI.1197-13.2013

Gilley, J., Coleman, M.P., 2010. Endogenous Nmnat2 Is an Essential Survival Factor for Maintenance of Healthy Axons. PLoS Biol. 8, e1000300. doi:10.1371/journal.pbio.1000300

Gilley, J., Jackson, O., Pipis, M., Estiar, M.A., Gan-Or, Z., Goutman, S.A., Harms, M.B., Kaye, J., Lima, L., Genomics, Q.S., Ravits, J., Rouleau, G.A., Züchner, S., Reilly, M.M., Coleman, M.P., 2021. Enrichment of SARM1 alleles encoding variants with constitutively hyperactive NADase in patients with ALS and other motor nerve disorders. medRxiv 2021.06.17.21258268. doi:10.1101/2021.06.17.21258268

Gong, B., Pan, Y., Vempati, P., Zhao, W., Knable, L., Ho, L., Wang, J., Sastre, M., Ono, K., Sauve, A.A., Pasinetti, G.M., 2013. Nicotinamide riboside restores cognition through an upregulation of proliferator-activated receptor-γ coactivator 1α regulated β-secretase 1 degradation and mitochondrial gene expression in Alzheimer’s mouse models. Neurobiol. Aging 34, 1581–1588. doi:10.1016/j.neurobiolaging.2012.12.005

Henninger, N., Bouley, J., Sikoglu, E.M., An, J., Moore, C.M., King, J.A., Bowser, R., Freeman, M.R., Brown, R.H., 2016. Attenuated traumatic axonal injury and improved functional outcome after traumatic brain injury in mice lacking Sarm1. Brain 139, 1094–1105. doi:10.1093/brain/aww001

Hikosaka, K., Ikutani, M., Shito, M., Kazuma, K., Gulshan, M., Nagai, Y., Takatsu, K., Konno, K., Tobe, K., Kanno, H., Nakagawa, T., 2014. Deficiency of nicotinamide mononucleotide adenylyltransferase 3 (nmnat3) causes hemolytic anemia by altering the glycolytic flow in mature erythrocytes. J. Biol. Chem. 289, 14796–14811. doi:10.1074/jbc.M114.554378

Horsefield, S., Burdett, H., Zhang, X., Manik, M.K., Shi, Y., Chen, J., Qi, T., Gilley, J., Lai, J.-S., Rank, M.X., Casey, L.W., Gu, W., Ericsson, D.J., Foley, G., Hughes, R.O., Bosanac, T., Itzstein von, M., Rathjen, J.P., Nanson, J.D., Boden, M., Dry, I.B., Williams, S.J., Staskawicz, B.J., Coleman, M.P., Ve, T., Dodds, P.N., Kobe, B., 2019. NAD+ cleavage activity by animal and plant TIR domains in cell death pathways. Science 365, 793–799. doi:10.1126/science.aax1911

Hughes, R.O., Bosanac, T., Mao, X., Engber, T.M., DiAntonio, A., Milbrandt, J., Devraj, R., Krauss, R., 2021. Small Molecule SARM1 Inhibitors Recapitulate the SARM1-/-Phenotype and Allow Recovery of a Metastable Pool of Axons Fated to Degenerate. Cell Reports 34, 108588. doi:10.1016/j.celrep.2020.108588

Jiang, Y., Liu, T., Lee, C.-H., Chang, Q., Yang, J., Zhang, Z., 2020. The NAD+-mediated self-inhibition mechanism of pro-neurodegenerative SARM1. Nature 588, 658–663. doi:10.1038/s41586-020-2862-z

Ko, K.W., Milbrandt, J., DiAntonio, A., 2020. SARM1 acts downstream of neuroinflammatory and necroptotic signaling to induce axon degeneration. The Journal of Cell Biology 219, 152. doi:10.1083/jcb.201912047

Krauss, R., Bosanac, T., Devraj, R., Engber, T., Hughes, R.O., 2020. Axons Matter: The Promise of Treating Neurodegenerative Disorders by Targeting SARM1-Mediated Axonal Degeneration. Trends Pharmacol. Sci. 41, 281–293. doi:10.1016/j.tips.2020.01.006

Li, W.H., Huang, K., Cai, Y., Wang, Q.W., Zhu, W.J., Hou, Y.N., Wang, S., Cao, S., Zhao, Z.Y., Xie, X.J., Du, Y., Lee, C.-S., Lee, H.C., Zhang, H., Zhao, Y.J., 2021. Permeant fluorescent probes visualize the activation of SARM1 and uncover an anti-neurodegenerative drug candidate. doi:10.7554/eLife.67381

Liu, H.-W., Smith, C.B., Schmidt, M.S., Cambronne, X.A., Cohen, M.S., Migaud, M.E., Brenner, C., Goodman, R.H., 2018. Pharmacological bypass of NAD+ salvage pathway protects neurons from chemotherapy-induced degeneration. Proc. Natl. Acad. Sci. U.S.A. 115, 10654–10659. doi:10.1073/pnas.1809392115

Loreto, A., Angeletti, C., Gu, W., Osborne, A., Nieuwenhuis, B., Gilley, J., Arthur-Farraj, P., Merlini, E., Amici, A., Luo, Z., Hartley-Tassell, L., Ve, T., Desrochers, L.M., Wang, Q., Kobe, B., Orsomando, G., Coleman, M.P., 2021. Potent activation of SARM1 by NMN analogue VMN underlies vacor neurotoxicity. bioRxiv 2020.09.18.304261. doi:10.1101/2020.09.18.304261

Marion, C.M., McDaniel, D.P., Armstrong, R.C., 2019. Sarm1 deletion reduces axon damage, demyelination, and white matter atrophy after experimental traumatic brain injury. Experimental Neurology 321, 113040. doi:10.1016/j.expneurol.2019.113040

Mayer, M., Meyer, B., 1999. Characterization of Ligand Binding by Saturation Transfer Difference NMR Spectroscopy. Angew Chem Int Ed Engl 38, 1784–1788. doi:10.1002/(SICI)1521-3773(19990614)38:12<1784::AID-ANIE1784>3.0.CO;2-Q

McCoy, A.J., Grosse-Kunstleve, R.W., Adams, P.D., Winn, M.D., Storoni, L.C., Read, R.J., 2007. Phaser crystallographic software. J Appl Crystallogr 40, 658–674. doi:10.1107/S0021889807021206

Osterloh, J.M., Yang, J., Rooney, T.M., Fox, A.N., Adalbert, R., Powell, E.H., Sheehan, A.E., Avery, M.A., Hackett, R., Logan, M.A., MacDonald, J.M., Ziegenfuss, J.S., Milde, S., Hou, Y.-J., Nathan, C., Ding, A., Brown, R.H., Conforti, L., Coleman, M., Tessier-Lavigne, M., Züchner, S., Freeman, M.R., 2012. dSarm/Sarm1 Is Required for Activation of an Injury-Induced Axon Death Pathway. Science 337, 481–484. doi:10.1126/science.1223899

Ozaki, E., Gibbons, L., Neto, N.G., Kenna, P., Carty, M., Humphries, M., Humphries, P., Campbell, M., Monaghan, M., Bowie, A., Doyle, S.L., 2020. SARM1 deficiency promotes rod and cone photoreceptor cell survival in a model of retinal degeneration. Life Sci Alliance 3, e201900618. doi:10.26508/lsa.201900618

Ratajczak, J., Joffraud, M., Trammell, S.A.J., Ras, R., Canela, N., Boutant, M., Kulkarni, S.S., Rodrigues, M., Redpath, P., Migaud, M.E., Auwerx, J., Yanes, O., Brenner, C., Cantó, C., 2016. NRK1 controls nicotinamide mononucleotide and nicotinamide riboside metabolism in mammalian cells. Nat Commun 7, 13103–12. doi:10.1038/ncomms13103

Sambeat, A., Ratajczak, J., Joffraud, M., Sanchez-Garcia, J.L., Giner, M.P., Valsesia, A., Giroud-Gerbetant, J., Valera-Alberni, M., Cercillieux, A., Boutant, M., Kulkarni, S.S., Moco, S., Cantó, C., 2019. Endogenous nicotinamide riboside metabolism protects against diet-induced liver damage. Nat Commun 10, 4291–11. doi:10.1038/s41467-019-12262-x

Sasaki, Y., Kakita, H., Kubota, S., Sene, A., Lee, T.J., Ban, N., Dong, Z., Lin, J.B., Boye, S.L., DiAntonio, A., Boye, S.E., Apte, R.S., Milbrandt, J., 2020. SARM1 depletion rescues NMNAT1-dependent photoreceptor cell death and retinal degeneration. Elife 9, 817. doi:10.7554/eLife.62027

Sasaki, Y., Milbrandt, J., 2010. Axonal degeneration is blocked by nicotinamide mononucleotide adenylyltransferase (Nmnat) protein transduction into transected axons. J. Biol. Chem. 285, 41211–41215. doi:10.1074/jbc.C110.193904

Sasaki, Y., Nakagawa, T., Mao, X., DiAntonio, A., Milbrandt, J., 2016. NMNAT1 inhibits axon degeneration via blockade of SARM1-mediated NAD+depletion. Elife 5, 1010. doi:10.7554/eLife.19749

Sasaki, Y., Vohra, B.P.S., Lund, F.E., Milbrandt, J., 2009. Nicotinamide mononucleotide adenylyl transferase-mediated axonal protection requires enzymatic activity but not increased levels of neuronal nicotinamide adenine dinucleotide. J. Neurosci. 29, 5525–5535. doi:10.1523/JNEUROSCI.5469-08.2009

Shen, C., Vohra, M., Zhang, P., Mao, X., Figley, M.D., Zhu, J., Sasaki, Y., Wu, H., DiAntonio, A., Milbrandt, J., 2021. Multiple domain interfaces mediate SARM1 autoinhibition. Proc. Natl. Acad. Sci. U.S.A. 118, e2023151118. doi:10.1073/pnas.2023151118

Sporny, M., Guez-Haddad, J., Khazma, T., Yaron, A., Dessau, M., Shkolnisky, Y., Mim, C., Isupov, M.N., Zalk, R., Hons, M., Opatowsky, Y., 2020. Structural basis for SARM1 inhibition and activation under energetic stress. Elife 9, W344. doi:10.7554/eLife.62021

Sporny, M., Guez-Haddad, J., Lebendiker, M., Ulisse, V., Volf, A., Mim, C., Isupov, M.N., Opatowsky, Y., 2019. Structural Evidence for an Octameric Ring Arrangement of SARM1. J Mol Biol 431, 3591–3605. doi:10.1016/j.jmb.2019.06.030

Summers, D.W., DiAntonio, A., Milbrandt, J., 2014. Mitochondrial dysfunction induces Sarm1-dependent cell death in sensory neurons. J. Neurosci. 34, 9338–9350. doi:10.1523/JNEUROSCI.0877-14.2014

Summers, D.W., Frey, E., Walker, L.J., Milbrandt, J., DiAntonio, A., 2019. DLK Activation Synergizes with Mitochondrial Dysfunction to Downregulate Axon Survival Factors and Promote SARM1-Dependent Axon Degeneration. Molecular Neurobiology 89, 449–13. doi:10.1007/s12035-019-01796-2

Turkiew, E., Falconer, D., Reed, N., Höke, A., 2017. Deletion of Sarm1 gene is neuroprotective in two models of peripheral neuropathy. Journal of the peripheral nervous system : JPNS 22, 162–171. doi:10.1111/jns.12219

Vonrhein, C., Flensburg, C., Keller, P., Sharff, A., Smart, O., Paciorek, W., Womack, T., Bricogne, G., 2011. Data processing and analysis with the autoPROC toolbox. Acta Crystallogr D Biol Crystallogr 67, 293–302. doi:10.1107/S0907444911007773

Wan, L., Essuman, K., Anderson, R.G., Sasaki, Y., Monteiro, F., Chung, E.-H., Osborne Nishimura, E., DiAntonio, A., Milbrandt, J., Dangl, J.L., Nishimura, M.T., 2019. TIR domains of plant immune receptors are NAD+-cleaving enzymes that promote cell death. Science 365, 799–803. doi:10.1126/science.aax1771

Wu, T., Zhu, J., Strickland, A., Ko, K.W., Sasaki, Y., Dingwall, C., Yamada, Y., Figley, M.D., Mao, X., Neiner, A., Bloom, J., DiAntonio, A., Milbrandt, J., 2021. Neurotoxins subvert the allosteric activation mechanism of SARM1 to induce neuronal loss. bioRxiv 2021.07.16.452689. doi:10.1101/2021.07.16.452689

Zhao, Z.Y., Xie, X.J., Li, W.H., Liu, J., Chen, Z., Zhang, B., Li, T., Li, S.L., Lu, J.G., Zhang, L., Zhang, L.-H., Xu, Z., Lee, H.C., Zhao, Y.J., 2019. A Cell-Permeant Mimetic of NMN Activates SARM1 to Produce Cyclic ADP-Ribose and Induce Non-apoptotic Cell Death. iScience 15, 452–466. doi:10.1016/j.isci.2019.05.001

